# The coevolution of learning schedules and teaching enhances cumulative knowledge and drives a teacher-innovator syndrome

**DOI:** 10.1101/2024.06.28.601211

**Authors:** Ludovic Maisonneuve, Laurent Lehmann, Charles Mullon

## Abstract

Natural selection shapes how individuals learn and acquire knowledge from their environment. Under the right conditions, this can lead to the evolution of learning schedules—how individuals allocate resources to acquire knowledge throughout their lifespan—that promote the accumulation of knowledge across generations (“cumulative knowledge” or “cumulative culture”). In spite of having been observed across multiple taxa, the role of parental teaching in this evolutionary process remains understudied. Using mathematical modelling, we show that learning schedules and parental teaching coevolve, resulting in greater time spent learning individually and innovating, as well as greater inter-generational transfer of knowledge from parent to offspring. These outcomes together enhance cumulative knowledge. Our analyses further reveal that within populations, selection typically favours an association between teaching and individual learning whereby some individuals innovate and teach within the family (“knowledge producers” with extensive knowledge), while others teach less and learn socially outside of the family (“knowledge scroungers” with less knowledge). Overall, our findings indicate that the coevolution of learning schedules and teaching promotes knowledge accumulation within and between generations, and favours diversity within and between populations in knowledge acquisition, possession and transmission.

## 1 Introduction

To improve their survival and reproduction, essentially all animals collect and use information about their surroundings [1]. This knowledge can be acquired from multiple sources as organisms can learn individually, e.g. through trial-and-error [2], as well as socially from conspecifics such as through imitation [3, 4]. Social transmission of knowledge itself can occur via many different pathways, which can usefully be classified as vertical (from parents), oblique (from other individuals of the parental generation), or horizontal (from peers of the same generation). How an individual allocates time and energy throughout its lifespan to the different activities and structures that underlie knowledge acquisition, we refer to as its learning schedule. Understanding selection on learning schedules is a challenging problem in life-history evolution owing to the multiple trade-offs that arise when allocating resources to different learning and other fitness-enhancing factors. This is compounded by the social dimension of the problem as the outcome of social learning depends on how much others know and consequently on others’ learning schedules.

Mathematical models have provided useful insights into how selection shapes learning schedules, particularly in identifying factors that favour individual against social learning in the form of oblique transmission, e.g. environmental variation [5–9], population structure and kin selection [10–12], or short memory [13]. One key result is that because social learning is often less costly than individual learning (e.g. less risky or more energetically efficient than trial-and-error), selection tends to favour a schedule whereby individuals first learn mostly socially and then increasingly through individual learning as they age and the amount of knowledge available socially decreases [14–17]. With such a schedule, an individual is able to build upon others’ knowledge within its own lifetime. This in turn opens the possibility for cumulative knowledge, i.e. an accumulation of knowledge across generations, such that eventually an individual can acquire more knowledge than it would through individual learning alone. Cumulative knowledge is considered to have played a central role in the evolution of hominin lineages [18], is at the heart of economic growth [19], and is thought to contribute to adaptation in other animals, such as crows [20], pigeons [21] and ungulates [22]. There has therefore been significant effort to identify the factors that favour the evolution of learning schedules conducive to cumulative knowledge, such as conditional learning [11, 23], low learning costs [14, 24], high social connectivity [25], knowledge-dependent learning [17], or interactions with relatives (via vertical learning [26, 27] or non vertical learning from other kin [10, 12, 28, 29]).

Evolutionary studies of cumulative knowledge have predominantly focused on the evolution of behaviours that are expressed by the learning individual (though see [30, 31]). Exemplar individuals from which learners obtain knowledge are typically considered as passive subjects with no agency over knowledge transmission. But from tandem running in ants that guide naive individuals toward new nests [32], through provisioning live prey to pups so they can practice capture in meerkats [33], to institutionalised instruction giving in hunter-gatherers [34], exemplar individuals across multiple taxa have been found to engage in activities that facilitate knowledge acquisition by the learner (see also [35–42] for other examples). Such behaviours facilitating knowledge acquisition by others constitute forms of teaching when they are costly to those performing them and are performed only in the presence of an observer [43]. We adopt this functional definition of animal teaching here, aligning with the view that teaching is an adaptation whose primary role is to enhance the learner’s ability to acquire knowledge [44–46]. There is nevertheless debate about whether the intention to assist the learner should not also be part of the definition of teaching [47, 48]; a perspective that maybe relevant but seems impossible to assess in non-human species. Regardless, teaching behaviours include increasing tolerance for intrusive observation, modifying the environment to support learning, or altering learner behaviour through rewards or penalties [44–46]. Because teaching is costly to its actor but beneficial to its recipient, it most readily evolves to be directed toward kin owing to inclusive fitness effects [30, 31, 49–53].

Teaching and learning schedules should co-evolve, coupled through a feedback via knowledge and its distribution in the population. This is because on the one hand, only certain learning schedules allow for kin selection to act on teaching. This is the case of schedules that involve vertical learning as these give the opportunity for parents to teach their offspring. On the other hand, because teaching increases the efficiency of knowledge transmission through certain pathways compared to others, it influences the learning schedule that will be favoured by selection. For instance if parents teach their offspring, it may be advantageous to spend more time and energy learning vertically at the expense of other learning pathways. Furthermore, any variation in teaching or in learning schedule leads to variation in knowledge in the population, which should in turn feed back on selection. Yet, the joint evolution of learning and teaching and its consequences are not well-understood, in part because the only study investigating this interplay does not permit for the accumulation of knowledge across generations [53].

Using mathematical modelling, we show here that the joint evolution of learning and parental teaching facilitates the establishment of cumulative knowledge and drives knowledge variation between families. This is due to a positive inter-generational feedback between parent and offspring leading to on average greater innovation within families, but also greater variance in innovation among families. Taken together, our findings indicate that teaching evolution has a wide potential to increase variation in knowledge within populations, and to enhance adaptive abilities in the many animal species that show parental care, including humans.

## 2 Model

We consider a large asexual population of haploids (the effects of mating and diploidy are discussed later). Each individual can collect and use knowledge, which we conceptualise as a quantitative variable that can be thought of as “embodied capital”; namely, a durable quantity accumulated by the individual during its lifespan that is used as input for its survival and reproduction (e.g., [15, 17, 54]). At each generation, the following life-cycle events unfold: (1) Parents produce a Poisson distributed number of offspring with mean depending on their knowledge. (2) Offspring accumulate knowledge through individual and social learning. (3) Parents die. (4) Offspring compete to become adults of the next generation, resulting in density-dependent regulation. We specify below how offspring learn (section 2.1) and how knowledge influences fitness (section 2.2).

### 2.1 Knowledge dynamics

We assume that during stage (2) of the life cycle offspring first learn from their parent (vertical learning) then from other adults (oblique learning) and finally by themselves (individual learning, Fig. 1a). During social learning (vertical or oblique), we envision learning as a foraging process for pieces of knowledge, which take time to search and handle. We assume that during this process, the rate at which an offspring encounters new pieces of knowledge is proportional to the difference between the knowledge that is available socially and the offspring’s current knowledge (Appendix A.1.1 for details and connection with previous models). A parent can expand energy into teaching their offspring to increase this rate during vertical learning, though expanding such energy is costly.

**Figure 1:**
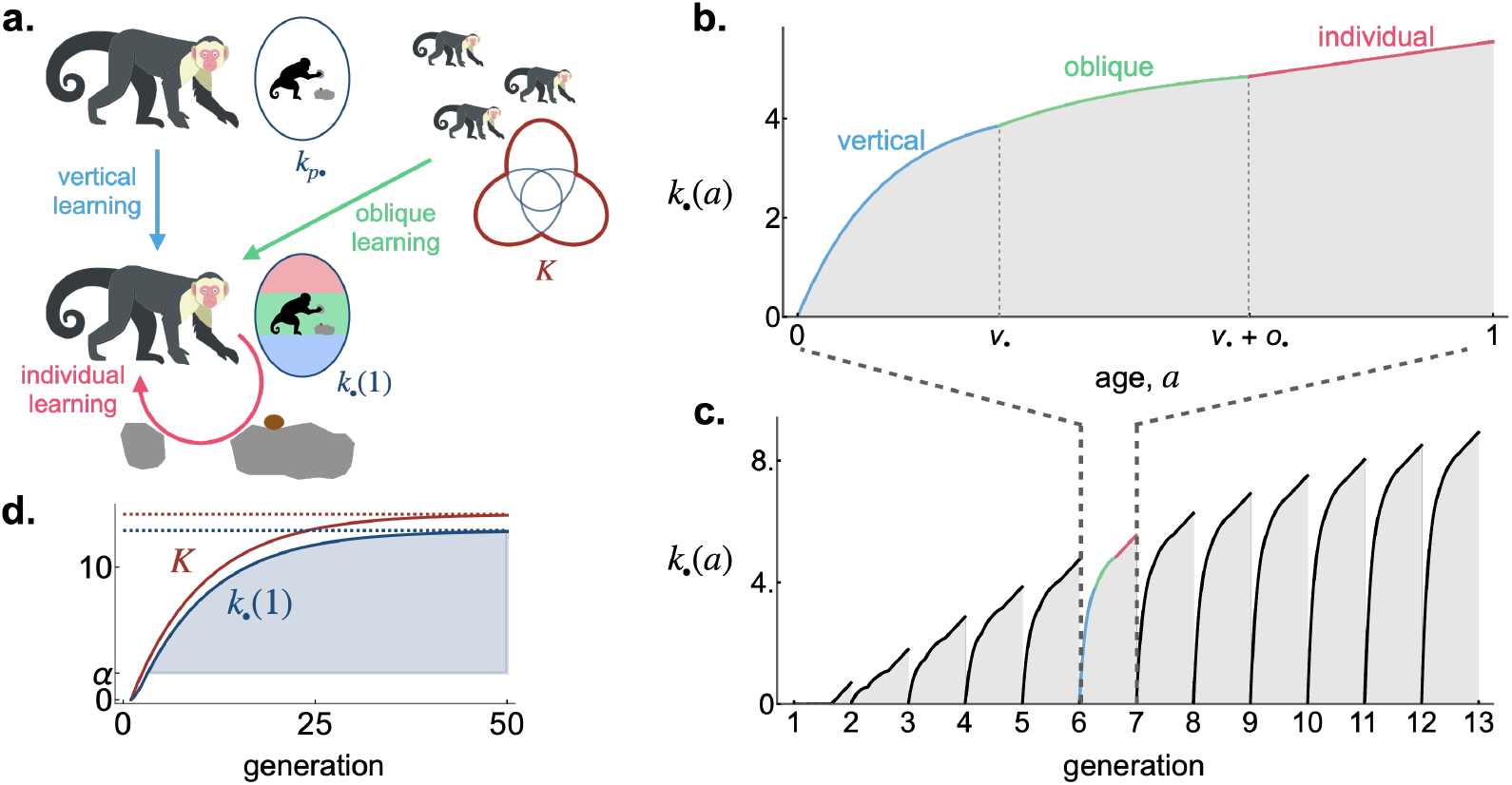
A model of cultural dynamics. **a**. A focal individual can obtain knowledge (e.g. the skill set to crack nuts open, denoted by *k*_*•*_(1) and represented here as the size of an oval set) by learning from three sources: (i) vertically from its parent (with knowledge *k*_p*•*_; blue arrow); obliquely from all adults in the population (with total knowledge *K*, which is represented as the size of the union in red of the knowledge of all adults to highlight that although each adult can possess unique pieces of knowledge, their knowledge can also overlap and thus be redundant; green arrow); and (iii) individually, when it generates its own knowledge (in pink). See main text section 2.1.1 for more details. **b**. Within-generation knowledge dynamics: individual knowledge *k*_*•*_(*a*) of a focal offspring against its age *a* (solution of eq. 1 with traits *v*_*•*_ *=* 0.29, *o*_*•*_ *=* 0.36, *τ*_*•*_ *=* 0.1 for the offspring and its parent; and parameters *β*_v0_ *=* 7, *β*_τ_ *=* 10, *β*_o_ *=* 4, *α =* 2, *ϵ =* 0.1, *ρ =* 0.5, *k*_p*•*_ *=* 4.75, *K =* 5.73). **c.-d**. Between-generation knowledge dynamics: Panel c shows individual knowledge *k*_*•*_(*a*) at each generation in a focal linage (same parameters as in panel b) in a population whose average trait values are *v =* 0.29, *o =* 0.36 and *τ =* 0.1 (using eqs. 1 and 3 starting with *K =* 0); Panel d shows individual knowledge *k*_*•*_(1) at the age of reproduction in the focal lineage (blue) and adults’ total knowledge *K* (red) at each generation (same parameters as in panel c; dashed lines indicate equilibrium values for knowledge, from eqs. A.19 and 6). Shaded area corresponds to cumulative knowledge (i.e. where *k*_*•*_(1) *> α*).

Knowledge that is obtained socially (via vertical or oblique learning) becomes obsolete at a rate *ϵ* from one generation to the next (Table 1 for a list of symbols). When *ϵ =* 0, knowledge acquired in one generation is as relevant in the next. In contrast, knowledge becomes irrelevant at each generation when *ϵ =* 1. The parameter *ϵ* can thus be thought of as a measure of environmental heterogeneity between generations.

**Table 1:** Key symbols and their definitions.

| Table 1: Key symbols and their definitions |  |
| --- | --- |
| Model parameters |  |
| $\epsilon$ | Rate of obsolescence of socially acquired knowledge (between-generation environmental heterogeneity) |
| $\rho$ | Probability that two individuals produce simultaneously different pieces of knowledge via individual learning (within-generation environmental heterogeneity) |
| $\alpha$ | Individual learning rate (eq. 1) |
| $\beta_o$ | Oblique transmission rate (eq. 1) |
| $\beta_{v0}$ | Baseline vertical transmission rate (eq. 2) |
| $\beta_\tau$ | Teaching efficiency (eq. 2) |
| $\gamma$ | Strength of density-dependent competition (eq. 4) |
| Variables for traits and their distribution |  |
| $\mathbf{x} = (v, o, \tau)$ | Mean trait values in the population: vertical learning time ($v$), oblique learning time ($o$), and teaching effort ($\tau$) |
| $\mathbf{G}$ | Variance/covariance matrix of trait values in the population |
| $\mathbf{x}_0^* = (v_0^*, o_0^*, 0)$ | Mean trait values favoured by selection when teaching is fixed at 0 |
| $\mathbf{x}^* = (v^*, o^*, \tau^*)$ | Mean trait values favoured by selection when all three traits co-evolve |
| $\mathbf{x}_\bullet = (v_\bullet, o_\bullet, \tau_\bullet)$ | Trait values of a focal individual |
| $\tau_{p\bullet}$ | Teaching effort of the focal's parent (eq. 1) |
| $\bar{v}_{o\bullet}$ | Average time spent learning vertically among the focal's offspring (eq. 4) |
| Variables for knowledge and its transmission |  |
| $k_\bullet(a)$ | Knowledge of a focal individual at age $a$ from birth ($a = 0$) to reproduction ($a = 1$, eq. 1) |
| $\beta_v(\tau_\bullet)$ | Vertical transmission rate when parent expresses teaching effort $\tau_\bullet$ (eq. 2) |
| $\xi_v(\mathbf{x}_\bullet)$ | Vertical inter-generational rate of knowledge transmission in a $\mathbf{x}_\bullet$-lineage (eq. 5) |
| $\xi_o(\mathbf{x}_\bullet)$ | Oblique inter-generational rate of knowledge transmission in a $\mathbf{x}_\bullet$-lineage (eq. 5) |
| $\hat{k}(\mathbf{x}_\bullet, \mathbf{x})$ | Equilibrium individual knowledge at the age of reproduction in a $\mathbf{x}_\bullet$-lineage in a population expressing $\mathbf{x}$ (eq. 5) |
| $\hat{k}(\mathbf{x}) = \hat{k}(\mathbf{x}, \mathbf{x})$ | Equilibrium individual knowledge in a population expressing $\mathbf{x}$ |
| $\hat{K}(\mathbf{x})$ | Equilibrium total knowledge in a population expressing $\mathbf{x}$ (eq. 6) |
| $\Lambda(\mathbf{x})$ | Inter-generational effects of teaching in a population expressing $\mathbf{x}$ (eq. 10) |
| Variables for fitness |  |
| $w(\mathbf{x}_\bullet, k_\bullet, n_o)$ | Fitness of a focal individual with traits $\mathbf{x}_\bullet$ and knowledge $k_\bullet$ when $n_o$ offspring are produced in population (eq. 4) |
| $\mathcal{C}_d(\tau)$ | Developmental cost of teaching effort $\tau$ (eq. 4) |
| $\mathcal{C}_e(\tau_e)$ | Cost of effective teaching effort $\tau_e$ (given by $\tau_e = \bar{v}_{o\bullet} \tau_\bullet$ for a focal individual, eq. 4) |

After learning, all individuals reproduce at the same age, which we normalise to 1. As a result, the amount of time spent learning vertically, obliquely and individually always sum to one, creating an inherent trade-off in the time apportioned to different learning activities. Individuals influence their learning schedule and the transmission of knowledge to their offspring via three quantitative traits: the amount of time *v* spent learning vertically; (ii) the amount of time *o* spent learning obliquely (so that 1 − *v* − *o* is spent learning individually); and (iii) the investment (or effort) *τ* into teaching when a parent. We specify next how knowledge dynamics depends on these traits, first within (section 2.1.1) and then between (section 2.1.2) generations.

#### 2.1.1 Within-generation knowledge dynamics

Consider an arbitrary generation and a focal offspring with traits (*v*_*•*_, *o*_*•*_, *τ*_*•*_) in that generation. Let us denote by *k*_*•*_(*a*) the amount of knowledge this offspring bears at age *a* ∈ [0, 1] (where *a =* 0 is birth and *a =* 1 is reproduction). Under the above assumptions (but ignoring information handling time for now, see Appendix A.1.2 for details), the rate d*k*_*•*_(*a*)/d*a* of change in this offspring’s knowledge is decomposed among three learning phases:

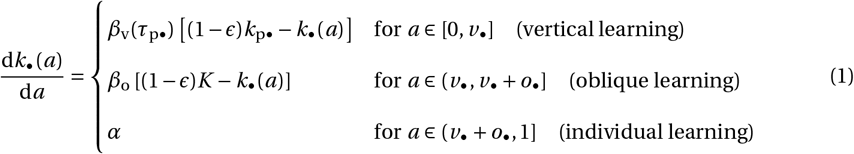

with initial condition *k*_*•*_(0) *=* 0 (Fig. 1b).

First, the offspring learns vertically for a time period of length *v*_*•*_ (first line of eq. 1), during which it encounters and assimilates knowledge at a rate that depends on: (i) a rate *β*_v_(*τ*_p*•*_) of vertical learning (detailed in eq. 2 below), and (ii) the difference [(1−*ϵ*)*k*_p*•*_ −*k*_*•*_(*a*)] between vertically available and the offspring’s current knowledge, where *k*_p*•*_ is the knowledge held by the offspring’s parent. We assume that *β*_v_(*τ*_p*•*_) increases with the teaching effort *τ*_p*•*_ of the offspring’s parent according to

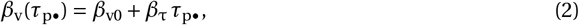

where *β*_v0_ is a baseline transmission rate that determines the vertical rate of learning in the absence of teaching (i.e. when *τ*_p*•*_ *=* 0), and *β*_τ_ is the conversion rate between teaching effort and learning rate, which can be thought of as a measure of teaching efficiency. Hence, an offspring can learn all the up-to-date knowledge of its parent (which is (1 − *ϵ*)*k*_p*•*_) when it spends a large amount of time learning vertically (*v*_*•*_ large), or when its parent invests much into teaching (*τ*_p*•*_ large).

In the second learning phase, the offspring learns obliquely from all the adults of the previous generation for a time period of length *o*_*•*_ (second line of eq. 1), during which it encounters and assimilates knowledge at a rate given by the product of: (i) the oblique transmission rate *β*_o_; and (ii) the difference [(1 −*ϵ*)*K* −*k*_*•*_(*a*)] between obliquely available and the offspring’s current knowledge, where *K* is a quantitative variable describing the total amount of distinct pieces of knowledge in the population (with each piece of knowledge counted only once, regardless of how many individuals share it). If the knowledge of each adult is conceived as a set, then this *K* can be thought of as the size of the union of all these sets in the population (see *K* in Fig. 1a). Because an adult may possess unique pieces of information that it has collected through individual learning, *K* is somewhere between the greatest individual knowledge among all adults and the sum of individual knowledge of all adults. We specify this *K* in the next section.

In a third and final phase, the offspring learns individually at a rate *α* for the remaining 1 − *v*_*•*_ − *o*_*•*_ time before reproduction (third line of eq. 1).

#### 2.1.2 Between-generations knowledge dynamics

As each offspring of a generation learns according to eq. (1), the total knowledge *K* in their generation increases, and this increase depends on whether they produce different or the same pieces of knowledge when they learn individually (e.g. developing independently the same skill). To capture this, we let *ρ* be the probability that two individuals from the same generation produce simultaneously different knowledge during individual learning. When *ρ =* 0, individuals always produce the same knowledge during individual learning so that *K* increases as much as the knowledge of just one individual. This may be because individuals encounter the same environmental features or stimuli, as would occur in a homogeneous environment. When *ρ =* 1, individuals always produce different and unique pieces of knowledge, thus systematically increasing total knowledge *K* during individual learning. This may occur because individuals encounter different environmental features or stimuli. The parameter *ρ* can be thought of as a measure of environmental heterogeneity within generations (while recall *ϵ* captures between-generations heterogeneity).

In Appendix A.2, we model the dynamics of *K* within and between generation from eq. (1) under the assumption that the population is large (see Fig. A.1 in the Appendix which shows that our approximation is excellent with as few as 100 individuals in the population, provided that individuals perform some social learning). For a population where the mean trait vector is ***x*** *=* (*v, o, τ*) and trait variance is small, we obtain that the change Δ*K* in the total knowledge over one generation is well approximated by

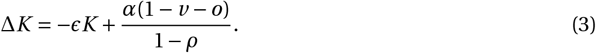

The term −*ϵK* is the loss of knowledge through obsolescence, and the second term is the new knowledge produced by individual learning in the population, which strongly depends on *ρ*. When *ρ* is close to 1 so that individuals produce different knowledge during individual learning, their collective contribution results in a substantial increase in total knowledge. By contrast when *ρ =* 0 so that individuals produce the same knowledge, the amount of new total knowledge is equal to the production of a single individual (which is *α*(1 − *v* −*o*)).

### 2.2 Individual fitness

We assume that knowledge enhances an individual’s reproductive success, but that teaching offspring is costly. Specifically, we assume that the effective fecundity, i.e. the expected number of offspring that reach adulthood of a focal individual with traits ***x***_*•*_ and knowledge *k*_*•*_(1) at the age of reproduction is given by

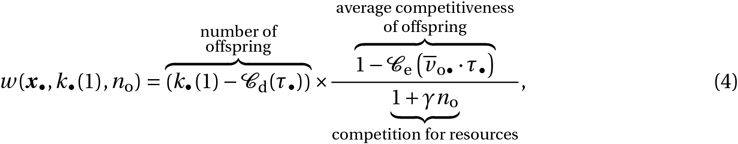

where 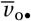 is the average time spent in vertical learning among the focal’s offspring, and *n*_o_ is the total number of offspring produced in the population. As indicated by the labels in eq. (4), effective fecundity results from three processes. Firstly, an individual produces a number of offspring that increases with the knowledge *k*_*•*_(1) it holds at the time of reproduction, but decreases with the developmental or metabolic cost 𝒞_d_(*τ*_*•*_) of having the ability to teach. This cost captures the notion that even if an individual does not effectively teach, e.g. if its offspring do not spend any time learning vertically, it still suffers a cost from developing the structures (e.g. investment into a specific cognitive module) that allow one to teach. Secondly, the competitiveness of offspring during density dependent competition decreases with the effective amount of teaching that their parent has done, reflecting that teaching may divert parents away from providing other resources. This effective cost of teaching, which we write as 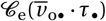, is assumed to increase with the amount of teaching conducted by the parent and the offspring vertical learning time according to the product 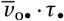. Thirdly and finally, offspring compete for resources following a Beverton-Holt model [55] of density-dependent competition where *γ* tunes the strength of this competition. The Beverton-Holt model, which can be considered as the discrete-time analogue of the continuous-time logistic equation for population growth, was chosen for its simplicity. Our model is therefore demographically explicit, allowing for the population size to fluctuate and to depend on how much knowledge individuals carry through their fecundity.

### 2.3 Evolutionary analyses

We assume that each trait, *v*, *o* and *τ*, evolves via rare mutations of small effects such that the distribution of traits in the population is approximately multi-variate Gaussian with small covariance. We track the evolution of the mean trait values ***x*** *=* (*v, o, τ*) and the variance-covariance matrix ***G*** of this distribution based on the approach detailed in [56] combined with that of [27]. This analysis allows us to determine whether selection is: (i) stabilizing, and in this case the trait associations shaped by mutation and selection; or (ii) disruptive, in which case polymorphism emerges in the population (i.e. the trait distribution goes from being unimodal to being bimodal); all this in the presence of interactions between relatives and inter-generational knowledge transmission in the population.

The gist of this quantitative genetics approach is to use the fact that when mutations are rare with small trait effects, evolutionary dynamics unfold more slowly than demographic and knowledge dynamics. Consequently, we can assume when studying evolutionary dynamics that knowledge within each lineage and population size have reached their equilibrium given current trait values. Selection can then be characterised from “lineage fitness”, which is the expected number of successful offspring produced by an individual randomly sampled from a lineage of individuals expressing traits ***x***_*•*_ *=* (*v*_*•*_, *o*_*•*_, *τ*_*•*_) – i.e. the “***x***_*•*_-lineage” consisting of an individual with traits ***x***_*•*_, its offspring, the offspring of the offspring, and so on – in a population that can be considered to be essentially monomorphic for mean trait values ***x***. Lineage fitness is given by 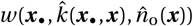 (with the function *w* given by eq. 4), where 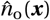 is the total number of offspring produced at demographic equilibrium in a population with mean trait values ***x***, and 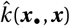 is the individual knowledge that an individual from the ***x***_*•*_-lineage carries at the age of reproduction (i.e. *k*_*•*_(1) at equilibrium of knowledge dynamics; Fig. 1cd for illustration of such dynamics). We show in Appendix A.3 that this equilibrium verifies

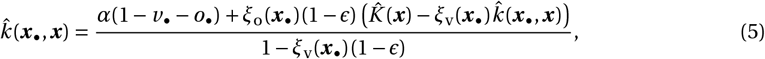

Where

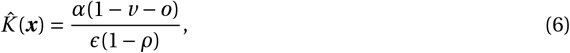

is the equilibrium total knowledge in the population (such that Δ*K =* 0 from eq. 3). The quantities 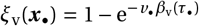 and 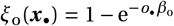 are the vertical and oblique inter-generational rates of knowledge transmission in the ***x***_*•*_-lineage; i.e. *ξ*_v_(***x***_*•*_) is the proportion of knowledge among that of a parent that is effectively learnt by its offspring (e.g. if a parent from the ***x***_*•*_-lineage has knowledge *k*_*•*_(1), it transmits *ξ*_v_(***x***_*•*_)(1 − *ϵ*)*k*_*•*_(1) to its offspring during vertical learning), and similarly, *ξ*_o_(***x***_*•*_) is the proportion of knowledge, among that can be transmitted obliquely, that is effectively learnt by the offspring. Eq. (5) decomposes the different factors that influence individual knowledge in a ***x***_*•*_- lineage. The first term in the numerator, *α*(1−*v*_*•*_−*o*_*•*_), is the knowledge that enters the lineage at each generation through individual learning while the second term is the knowledge entering through oblique learning. The denominator of eq. (5), meanwhile, captures the trans-generational accumulation of knowledge within the lineage due to vertical learning. To see this, we can unpack the reciprocal of the denominator as 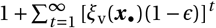, which shows the accumulation of non-obsolete knowledge that is transmitted vertically within the lineage across all generations. Altogether, eq. (5) thus states that the equilibrium knowledge within a lineage is determined by a balance between the knowledge continuously gained—through production within the lineage and acquisition from adults outside the lineage—and the knowledge continuously lost due to obsolescence and imperfect transmission between generations. We refer readers to eq. (A.19) in the Appendix for the explicit expres-sion of 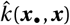, and to eq. (11) in the Methods for 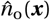, which depends on the equilibrium individual knowledge 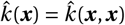 in a population expressing ***x***.

Since traits are genetically determined, selection in our model acts at the level of the gene, shaping the trait distribution within the population. Gene-level selection necessarily incorporates kin selection effects, which here arise because the knowledge of an individual—and consequently its fitness—is influenced not just by its own traits but also by those of its ancestors through vertical knowledge transmission [27]. For further information on our approach, see the Methods sections and Appendix B. Our evolutionary analyses are detailed in Appendix C and our main results are summarised below.

## 3 Results

### 3.1 Where vertical, oblique, and individual learning combine

We first investigate the evolution of learning schedules in the absence of teaching; that is, we fix *τ =* 0 (so that the vertical learning efficiency is just *β*_v0_, eq. 2) and assume that the situation is such that at least some level of social learning is favoured by selection when there is no teaching (requiring that eq. 16 is satisfied). We study the mean trait values, 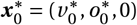, that evolve under directional selection (i.e. the convergence stable learning schedule) in a population where there is initially only individual learning. We are particularly interested in delineating the conditions where vertical, oblique, and individual learning are all performed at the evolutionary equilibrium, i.e. where 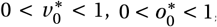, and 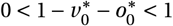 all hold. Because an individual cannot reproduce without knowledge in our model (see eq. 4), selection always favours some level of individual learning. The question is therefore when does selection also favours social learning, in both vertical and oblique forms.

We show that in the absence of teaching, an evolutionary equilibrium 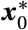 that combines all forms of learning is such that

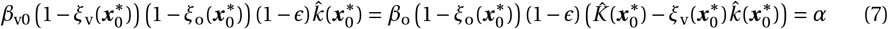

(Appendix C.3.1 for derivation). The left hand side of eq. (7) is the marginal effect of an increase in the time learning vertically by an individual on its amount of knowledge at reproduction (in a population expressing 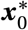). Likewise, the term in between equality signs is the marginal effect of an increase in the time spent learning obliquely on the amount of knowledge at reproduction (see Appendix C.3.2 for details). Finally, the right hand side of eq. (7) is the marginal effect of an increase in the time spent learning individually. Eq. (7) thus says that at 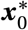, the marginal benefits of vertical, oblique, and individual learning on individual knowledge must balance each other out.

For this balance to be achieved, we show that it is necessary for

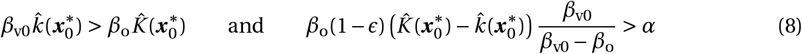

to hold (Appendix C.3.3 for derivation). When the first condition of (8) holds, a newborn offspring obtains knowledge at a rate that is faster via vertical 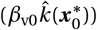 than via oblique learning 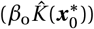. It is intuitive that selection in that case favours offspring that start learning from their parent rather than from the rest of the population. Because the total knowledge 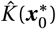 that is available through oblique learning is necessarily greater than the amount 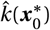 through vertical learning, the condition 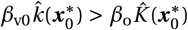 requires that the baseline efficiency of vertical learning is greater than the efficiency of oblique learning, *β*_v0_ *> β*_o_. This may for instance occur when strong bonds among parent and offspring entail that imitating a parent occurs more easily than imitating a non-kin. If, however, the first condition of (8) is not met, then selection necessarily leads individuals to skip vertical learning and start their lifespan with oblique learning (Fig. S1a.).

To understand the second condition of (8), it is useful to recognise that 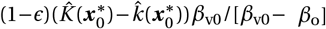 is the amount of knowledge available from oblique learning at the end of the vertical learning phase (see eq. C.18 in the Appendix). Therefore, when the second condition of (8) holds an individual after having learnt vertically, obtains knowledge at a faster rate via oblique learning (left hand side) than via individual learning (right hand side). It makes sense that selection then favours the evolution of oblique learning. If however the second condition of (8) does not hold, selection then favours skipping oblique learning (Fig. S1b.). This tends to happen when the amount of knowledge available from one’s parent is comparable to the knowledge available from the general population (i.e. where 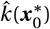 and 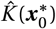 are similar such that the left hand side is small).

In sum, eq. (7) and conditions (8) reveal that selection favours the evolution of a combination of vertical, oblique, and individual learning, where in early life, offspring learn more efficiently from their parent than from unrelated individuals (*β*_v0_ *> β*_o_). This allows vertical learning to be favoured in spite of the fact that the amount of knowledge accessible outside the family is always greater than within (i.e. 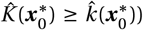. Once an offspring has acquired sufficient knowledge from its parent, however, it becomes more attractive to shift to oblique learning in order to tap into this wider pool of total knowledge (at age 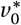, Fig. 2a.). In turn, as the difference between individual and total knowledge shrinks, there comes a time when the offspring can acquire knowledge more rapidly through individual learning (at age 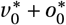, Fig. 2a.). At this point, the offspring thus makes a second shift in its learning schedule to produce its own knowledge, which, provided it has not become obsolete by then, can then be passed onto the next generation.

**Figure 2:**
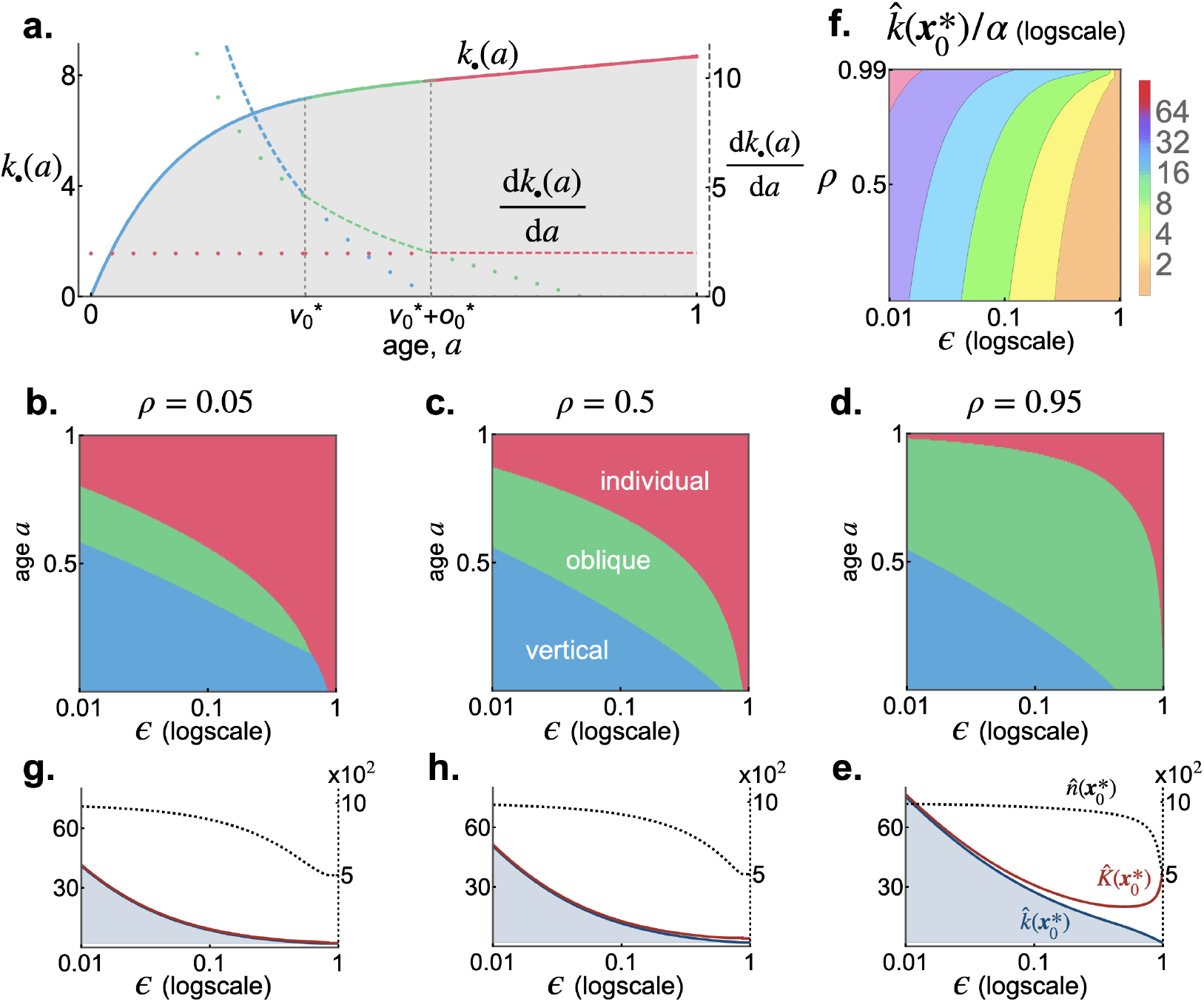
Learning schedule evolution in the absence of teaching. **a**. Individual knowledge *k*_*•*_(*a*) (full lines) and learning rate d*k*_*•*_(*a*)/d*a* (dashed lines) of an individual expressing the evolutionary equilibrium 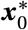 according to its age *a* with *τ =* 0 fixed (from eq. 1 and its solution, with *ϵ =* 0.1, *ρ =* 0.05, other parameters: same as Fig. 1b., see Appendix C.3.4 for procedure to find 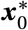). Dotted lines indicate the learning rates if the individual was performing vertical, oblique, and individual learning (in blue, green, and red, respectively) instead of the evolutionary equilibrium. This panel therefore shows that at 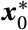, individuals switch to the learning mode that affords the greatest learning rate during their lifetime (dashed line). **b-d**. Learning schedule (y-axis) at evolutionary equilibrium 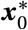 against *ϵ* (x-axis) for different values of *ρ* (other parameters: same as a.). Blue, green, and pink areas represent time spent performing vertical, oblique, and individual learning, respectively. **g**., **h**., **e**. Individual knowledge 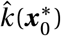 (blue) at the age of reproducing, total knowledge 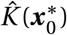 (red), and population size 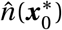 (black dashed) at 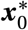 corresponding to panels b-d (left axis gives scale of knowledge, and right axis gives scale of population size, with *γ =* 10^−3^). Shaded area corresponds to cumulative knowledge (i.e. where 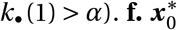 according to *ϵ* and *ρ* (other parameters: same as a.).

To gain more quantitative insights into how selection shapes learning, we determined numerically the trait values 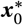 favoured by selection for a wide range of values for *ϵ* and *ρ* (Appendix C.3.4 for details on approach). Our results are summarised in Fig. 2. We find as expected that social learning (either vertical or oblique) is favoured relative to individual learning when *ϵ* is small since this means that the knowledge of previous generations remains up-to-date (Fig. 2b.-d.). Our analyses also reveal that vertical learning is favoured compared to oblique learning when *ρ* is small, so when individual learning tends to produce the same knowledge among individuals (Fig. 2b.). This is because the difference between knowledge within and outside the family is small in this case (i.e. 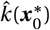 and 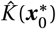 are similar, Fig. 2g.). As a consequence, the relative advantage of oblique learning is also small. In fact, this relative advantage becomes so small where *ρ* is small and *ϵ* sufficiently large, that selection favours skipping oblique learning altogether and instead leads to the evolution of a learning schedule that consists of vertical followed immediately by individual learning (e.g. Fig. 2g. with *ϵ* around 0.8 and Fig. S1b.)

We investigated the effect of learning evolution on cumulative knowledge by comparing individual knowledge 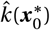 at 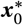, and *α* which corresponds to the knowledge an individual would obtain from learning individually only (i.e. where *v*_*•*_ *+o*_*•*_ *=* 0). Cumulative knowledge occurs where 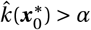. We find that cumulative knowledge is greatest where *ρ* is large and *ϵ* is small. Large *ρ* means that indi-viduals collectively can create large amounts of total knowledge through individual learning, while small *ϵ* entails that this knowledge remains up-to-date across many generations and can therefore accumulate within lineages through social learning. This accumulation readily results with individuals carrying knowledge in quantities one order of magnitude greater than through individual learning alone (Fig. 2f.; especially when oblique transmission rate *β*_o_ is high Fig. S2a.). This is accompanied by a corresponding increase in population size as fecundity is linked to knowledge in our model (Fig. 2g., h., e.).

### 3.2 The evolution of teaching boosts cumulative knowledge

Against a backdrop where teaching is absent, we investigate next whether selection favours the emergence of teaching. We show that in a population at 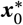, teaching evolves when

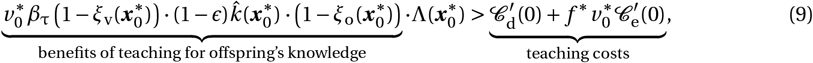

where 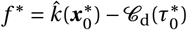 is the fecundity in a population at 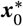, and

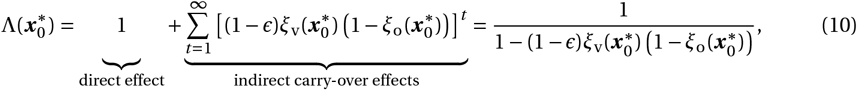

captures the inter-generational effects of teaching (Appendix C.4.1 for derivation). The right hand side of inequality (9) quantifies the marginal individual costs of teaching, composed of the developmental 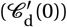 and effective 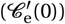 costs (a ′ denotes the derivative). Meanwhile, the left hand side of inequality (9) captures the marginal benefit of teaching, which is given by the product of two factors: (i) the positive effects of teaching by a parent on the knowledge of its surviving offspring (underbraced term in left hand side of inequality 9), which depend on the effects of teaching on the vertical transmission rate 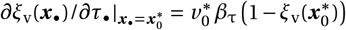 (Appendix C.4.2 for more details); and (ii) 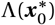, which captures the inter-generational effects of teaching since teaching under vertical transmission not only improves the knowledge of immediate descendants (“direct effect” in eq. 10), but also of the descendants of these and so on (“indirect carry-over effects” in eq. 10). These indirect effects consist in the effective share of knowledge, 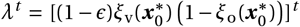, that an individual transmits to its descendants living *t* generations later, summed over all *t* (Fig. 3a for illustration). Such inter-generational effects are strongest – i.e. 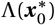 is largest and the potential impact of teaching on the knowledge of one’s descendants is greatest – when the rates of knowledge transmission between generations is large via vertical (large 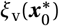) but small via oblique (small 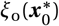) routes (Fig. S3a.). Accordingly, selection for the emergence of teaching is strongest under conditions that favour the evolution of vertical but not oblique learning (such that 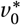 is large and 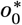 is small). This occurs when *ρ* is low and *ϵ* is intermediate (Fig. 3b. and Fig. S3b.).

**Figure 3:**
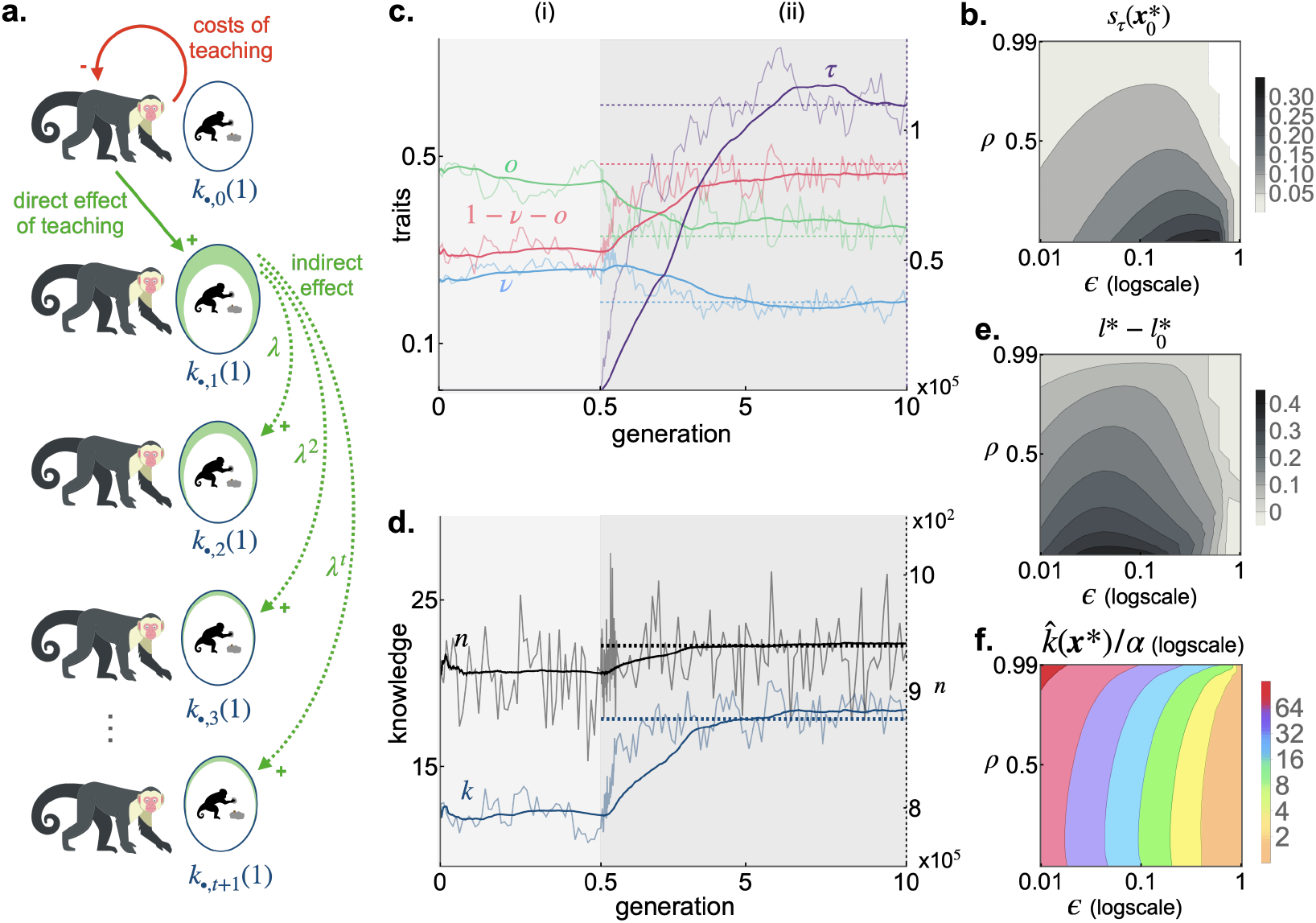
The coevolution of learning schedules and teaching. **a**. Direct costs and indirect inter-generational benefits of teaching (see main text section 3.2 for interpretation). **b**. The strength of selection for the emergence of teaching 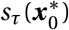 according to *ϵ* and *ρ* (with 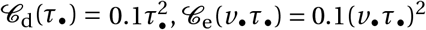, 𝒞 (*v*_*•*_*τ*_*•*_) *=* 0.1(*v*_*•*_*τ*_*•*_)^2^; other parameters: same as Fig. 1b.). **c-d**. The evolution of teaching and its impact on learning schedules (panel c; left axis gives scale of *v*, *o* and 1−*v* −*o*, and right axis gives scale of *τ*), and on knowledge and population size (panel d; left axis gives scale of knowledge, and right axis gives scale of population size). Thin lines show results of individual based simulations starting with a population monomorphic for 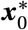 (Appendix C.5 for details, with *γ =* 10^−3^, *ϵ =* 0.1, *ρ =* 0; other parameters: same as b); Thick lines shows the moving mean of these (window length: 25’000); Dashed lines show evolutionary equilibrium obtained from mathematical model, i.e. show ***x***^*^ in c, and 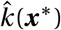 and 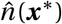 in d (Appendix C.4.3 for procedure to compute ***x***^*^). During period (i), *τ =* 0 is fixed so that traits stabilise for 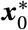. During period (ii), we allow mutations to affect *τ*, leading to the evolution of teaching (in purple), resulting in an increase in individual learning (in red). This in turn leads to a boost in knowledge (in blue) and thus population size (in black), though this latter is moderate due to competition for resources (eq. 4). **e-f**. Effect of the evolution of teaching on individual learning (in panel e, where *l*^*^ *=* 1 − *v*^*^ − *o*^*^ and 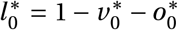) and cumulative knowledge (in panel f). These show that the evolution of teaching leads to greater time spent learning individually and greater cumulative knowledge. Parameters: same as b.

When teaching emerges, this affects the rate of vertical transmission and therefore selection on learning traits, whose evolution in turn feeds back on teaching. To investigate the evolutionary consequences of these feedbacks, we computed numerically the trait values ***x***^*^ *=* (*v*^*^, *o*^*^, *τ*^*^) favoured by directional selection, starting with a population expressing 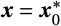 (Appendix C.4.3 for details). This analysis reveals that the evolution of teaching is typically accompanied with an increase in individual learning (i.e. 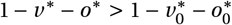, e.g. Fig. 3c. and 3e.). This occurs because teaching allows one’s offspring to rapidly acquire knowledge early on via vertical learning. An offspring thus quickly reaches the critical level of knowledge where it becomes more productive to shift to individual learning (e.g. Fig. S4). Because the evolution of teaching results both in: (i) greater individual learning, which leads to the creation of new knowledge; and (ii) an increase in vertical transmission rate, which ensures efficient inter-generational transmission of this new knowledge; the evolution of teaching causes an overall boost in cumulative knowledge (Fig. 3c. d. and Fig. S3c.) and thus in fecundity and population size (Fig. S3d.). Cumulative knowledge at ***x***^*^ is especially high when teaching is efficient, i.e. *β*_τ_ is large (Fig. S2b.).

The above results ignore the time it takes to handle information during social learning (eq. 1). We explore the effects of handling time on ***x***^*^ in Appendix D. Our analyses reveal that handling time leads to the evolution of lower teaching effort. This is because handling time reduces the rate of knowledge acquisition during vertical learning, thereby decreasing the benefit of teaching (Fig. D.1a). When handling time becomes sufficiently large, social learning becomes so inefficient that natural selection favours skipping both vertical and oblique learning, which prevents the evolution of teaching altogether (Fig. D.1a,b). Because handling time limits the efficiency of knowledge transmission, it also curtails cumulative knowledge and population size (Fig. D.1c).

### 3.3 A teacher-innovator syndrome with knowledge producers and scroungers

So far, our analyses have focused on the mean trait values ***x***^*^ that evolve under directional selection. We now investigate how selection shapes the distribution of trait values around these means. Specifically, we determine whether selection is stabilising or disruptive in a population at ***x***^*^, and in the former case, the between-traits correlations that evolve under mutation-selection balance (see ‘Dynamics of the Trait Variance-Covariance Matrix ***G***’ in the Methods for details).

We find that for most parameters we examined, selection is stabilising so that the phenotypic distribution remains unimodally distributed with mean ***x***^*^ (white region in Fig 4a.). In addition, selection typically shapes between-traits correlations such that individuals who teach more than others as adults, also tend to perform less of vertical and more individual learning as offspring (Fig. S5). This trait association, also observed in individual-based simulations (Fig 4b.), forms a suite of correlated behaviours known as a behavioural syndrome [57, 58]. We refer to this particular syndrome as a ‘teacher-innovator syndrome’. It emerges because in lineages where adults put greater effort into teaching, vertical learning is more efficient and allows offspring to spend less time learning vertically (unless *ϵ* is high, Fig. S5). Overall, offspring enjoy a boost in knowledge acquisition via vertical transmission. This boost in turn allows them to allocate more time to individual learning and therefore to innovation and the production of new knowledge. As a result, individuals from these lineages tend to carry more knowledge (Fig. S6a.).

**Figure 4:**
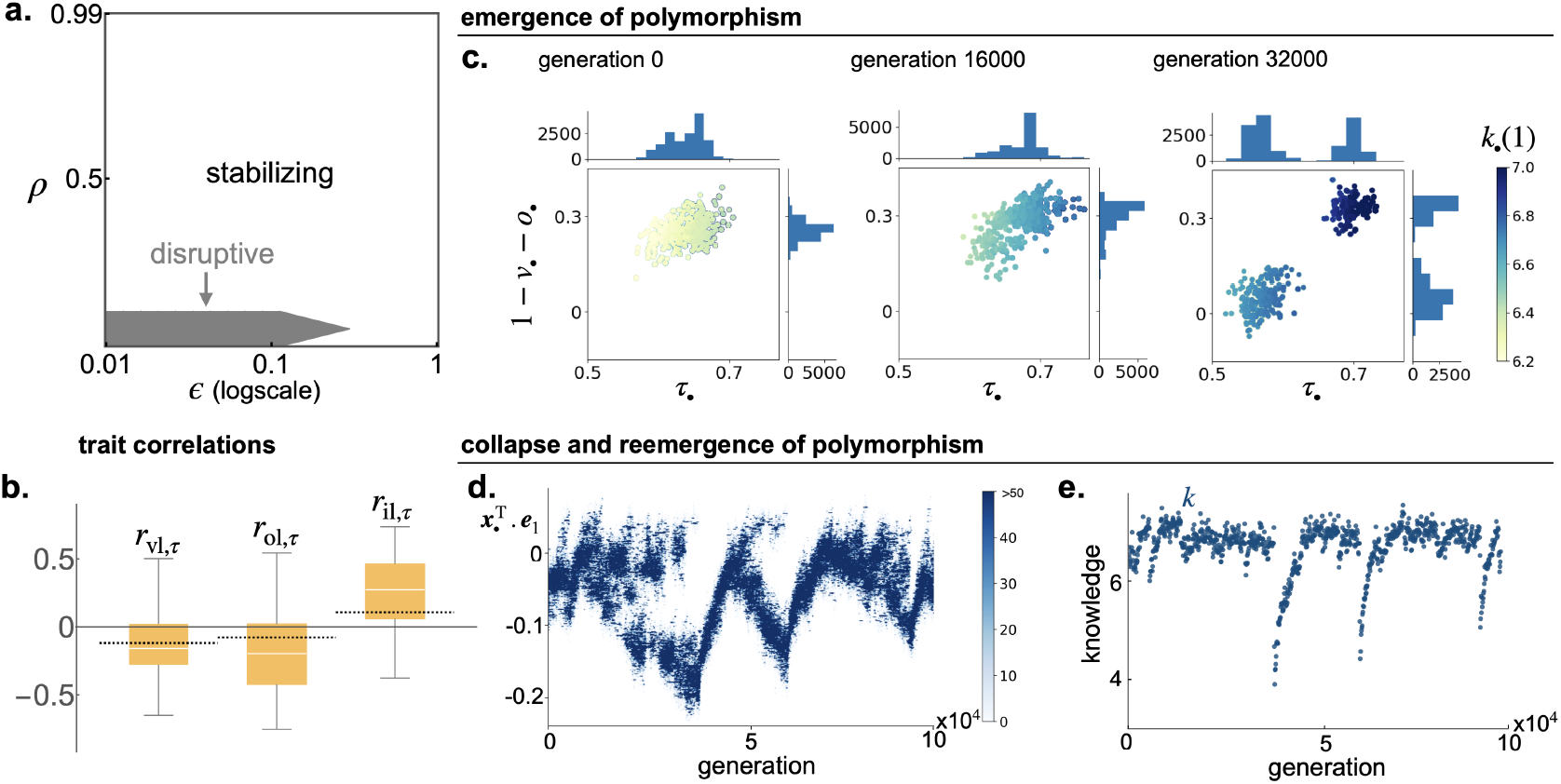
The evolution of a teacher-innovator syndrome. **a**. Space of parameters where selection is stabilising – so that the trait distribution in population remains unimodal around ***x***^*^ – or disruptive, favouring an increase in trait variance and the emergence of polymorphism (Parameters: same as in Fig 3b.). **b**. Correlation at selection-mutation balance when selection is stabilising between teaching effort and: (i) vertical learning *r*_vl,*τ*_; (ii) oblique learning *r*_ol,*τ*_; and (iii) individual learning *r*_il,*τ*_. Dashed black lines show equilibrium from mathematical model (from eq. B.10 in Appendix). Box plots show distribution over 80’000 generations from individual-based simulations (Appendix C.5 for details). With *ρ =* 0.5 and *γ =* 5 *×* 10^−5^, other parameters: same as in Fig 3. **c**. Snapshots of trait distribution from individual based simulations under disruptive selection (with parameters *β*_v0_ *=* 2, *β*_*τ*_ *=* 8, *β*_o_ *=* 1, *α =* 1, *ϵ =* 0.05, *ρ =* 0.05, 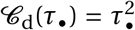, 𝒞_e_(*v*_*•*_*τ*_*•*_) *=* 0.5(*v*_*•*_*τ*_*•*_)^2^, *γ =* 5 *×* 10^−5^). This shows the emergence of two morphs: (i) some that spend more time learning individually as offspring and put more effort into teaching as parent; (ii) and others that spend less time learning individually as offspring and put less effort into teaching as parent. The former carry more knowledge than the latter at the age of reproduction *k*_*•*_(1) (see colour legend). **d-e**. Long term trait evolution under disruptive selection and its impact on knowledge (from same individual based simulations as in panel c). Panel d shows the trait distribution projected onto the subspace generated by the eigenvector ***e***_1_ of the Hessian matrix (eq. 15), which associated with the matrix’s leading eigenvalue (***e***_1_ determines the direction of disruptive selection in trait space [96, 97]). Panel e shows the average individual knowledge in the population at the age of reproduction (shown for every 100 generation). These two panels reveal the cycles of emergence and collapse of polymorphism, which generate cycles in knowledge (and thus also in population size, see Fig. S8).

By contrast, in lineages where there is less teaching, offspring spend less time on individual learning but more on oblique learning (Fig. S5). These offspring therefore tend to absorb knowledge from the population at large rather than from their parent. However, because social transmission of knowledge is weaker where there is less teaching, individuals who belong to such lineages tend to carry less knowledge (Fig. S6a.). In spite of this, these individuals nevertheless have fecundity that is similar to others (Fig. S6b.) since they avoid some of the costs that come with teaching.

Selection to associate teaching and individual learning requires *ϵ* and *ρ* to be low enough, i.e. low enough between- and within-generation environmental heterogeneity (Fig. S5). This is because the combination of increased teaching in parents, together with extended vertical and individual learning in offspring, has synergistic effects on knowledge accumulation across generations within lineages that are exacerbated in such environments. In fact, for some especially small values of *ϵ* and *ρ* (gray region in Fig 4a.), these synergistic effects can be so strong that selection becomes disruptive and leads to an increase of phenotypic variance. Individual-based simulations reveal that disruptive selection leads to the emergence of two discrete morphs: (i) “knowledge producers” that commit more efforts into teaching and more time into individual learning, and thereby contribute to the majority of knowledge production in the population; and (ii) “knowledge scroungers” that commit less efforts into teaching and more time into oblique learning, and thus benefit from the knowledge produced by the other morph (Fig. 4c and Fig. S7). Similar to classical producer-scrounger systems [59–63], scroungers depend on producers to acquire resources, which here is knowledge. This dependency leads to negative frequency-dependent selection that promotes polymorphism: scrounging is favoured by selection when scroungers are rare but disfavoured as their frequency increases.

Although both knowledge producers and scroungers are initially maintained under negative frequency dependent selection, individual-based simulations reveal that producers become so rare that they eventually go extinct stochastically (i.e. they become so few that by chance, none reproduce, Fig. 4d.). Here, frequency dependent selection favours a high frequency of knowledge scroungers and a correspondingly low frequency of producers (Appendix C.6 for details). This is because total knowledge *K*, which is the resource that is being produced and scrounged in our model, is only weakly impacted by morph frequencies. Two factors contribute to such weak effects of morph frequencies. First, since polymorphism emerges in environments where individuals tend to produce the same knowledge (i.e. low *ρ*), total knowledge production depends weakly on the number of producers in the population (unless there are very few). Second, the nature of knowledge, which here is a resource that can be considered a public good (i.e. non-excludable and non-rivalrous), means that having more scroungers in the population has no direct impact on total knowledge, i.e. scroungers do not “consume” the resource. This contrasts with classical producer-scrounger systems where the resource is typically rivalrous (i.e. a common-pool resource, e.g. food [59–63]).

Once the producer morph goes extinct, the population becomes monomorphic for the scrounger morph (Fig. 4d.). This extinction causes a major and sudden drop in knowledge since it is no longer being produced in the population (Fig. 4e.). Accordingly, the loss of polymorphism is accompanied by a drop in population size of similar intensity (Fig. S8). As scroungers can no longer benefit from the knowledge produced by others, selection acts against them and leads the population back towards ***x***^*^, resulting in an increase in both individual and total knowledge. However, once the population expresses ***x***^*^, selection becomes disruptive again, favouring the emergence of knowledge scroungers and producers, thus starting the cycle of evolutionary and cultural dynamics all over again. Over long time scales, we therefore observe the periodic emergence and collapse of a polymorphism that underlies cycles of knowledge and population size (Fig. 4d., Fig. S8).

## 4 Discussion

Our model demonstrates that the joint evolution of learning schedules and teaching can lead to a significant increase in the amount of knowledge that accumulates across generations in a population (Fig. 3d). Previous discussions of the effects of teaching on cumulative knowledge and/or culture revolved around its positive impact on transmitting past innovations [30, 31, 64, 65]. Here we have shown an additional path through which teaching improves cumulative knowledge: the evolution of teaching modulates the trade-offs among different learning activities and thus impacts how se-lection shapes learning schedules. Specifically, by enabling individuals to acquire socially available knowledge more rapidly, teaching favours a reduction in the time individuals spend on social, especially oblique, learning (Fig. 3c). This in turn allows individuals to allocate more time to individual learning and innovations, which produces new knowledge and thereby boosts cumulative knowledge (Fig. 3d).

Our results also indicate that the nature of the environment is important to the joint evolution of learning schedules and teaching. Specifically, the evolution of both vertical and oblique learning is favoured in environments where social knowledge remains up-to-date between generations (i.e., low *ϵ*), such as when between-generations environmental heterogeneity is low (Fig. 2). This finding aligns with much previous theory showing that oblique learning is selected over individual learning in temporally stable environments [5–8].

Additionally, the environment influences the relative benefits of vertical versus oblique learning. While vertical transmission may occur at a higher rate than oblique transmission due to the close parent-offspring relationship, oblique learning provides access to a greater amount and diversity of knowledge because it involves learning from a wider range of individuals outside the family. A previous model suggests that when the environment changes between generations, oblique learning is favoured by selection over vertical learning [66]. In this model, knowledge is discrete (i.e., individuals either have or do not have adaptive knowledge), and individuals can engage in only one type of learning: vertical, oblique, or individual. Consequently, oblique learners can acquire up-to-date knowledge from individual learners when the environment changes whereas vertical learners are likely to receive outdated knowledge from their parents. In our model where knowledge is quantitative and individuals can potentially perform all three learning types in their lifetime, the key factor determining whether selection favours oblique or vertical learning is the extent to which different individuals acquire different pieces of knowledge through individual learning — captured by our parameter *ρ*. Oblique learning evolves most readily when this *ρ* is large (Fig. 2b-d) so that the amount of knowledge that is accessible outside of the family is larger than within. This is likely to obtain in environments that show within-generation heterogeneity, such that different individuals are more likely to encounter different features or stimuli. In contrast, where within-generation environmental heterogeneity is low (small *ρ*), one expects less variation in knowledge among families so that selection tends to promote vertical learning. This in turn sets the stage for the evolution of teaching and an increase in cumulative knowledge.

Our study also provides insights into how selection can influence variation in learning and teaching within populations. We have shown that the trait distribution in the population at mutation-selection balance (and -drift in simulations) typically displays a correlation whereby individuals that invest more resources into teaching also tend to create more knowledge though individual learning (Fig. 4). This results from the synergistic inclusive fitness effects of the traits that modulate learning and teaching: lineages of individuals who teach more can spend less time on social learning and more time on individual learning thereby innovating more. Individuals from these lineages typically carry greater knowledge than others, which increases their fecundity and/or survival; an increase, however, that is offset by teaching costs. This in turn opens a niche for individuals with a lower teaching load as these avoid teaching costs, but can still benefit from the knowledge of producer lineages through oblique learning. Where the synergy is strong enough, this evolutionary process can result in the emergence of two discrete morphs although these two morphs are not maintained in equilibrium as producer lineages become increasingly rare until they become extinct. This contrasts with the polymorphism described in [50], where it is found that in a population where oblique learning occurs from a randomly sampled individual in the population, teachers and non-teachers can stably coexist. In our model where oblique learning occurs from any other individual in the population that carries more units of knowledge, producer lineages have a greater disadvantage as they can be more easily exploited, leading to their demise. Once producers go extinct in our model, this restarts the process of polymorphism emergence and collapse, resulting in cycles of boom and bust of knowledge (Fig. 4d-e) and consequently of population size (Fig. S8).

The boom-and-bust cycles we report could stabilise if producers were to evolve strategies that limit their exploitation by scroungers, such as masking their knowledge to prevent transmission. A real-world example of this behaviour is observed in baboons (*Papio ursinus*), where individuals deliberately wait for others to leave before approaching a food source, thus hiding the location from potential competitors [67]. When knowledge pertains to a consumable resource—such as the location of food that can be depleted—this kind of masking becomes especially advantageous as it prevents others from accessing and depleting the resource. In these cases, knowledge itself becomes consumable, losing its value once the resource it pertains to has been exploited. This stands in contrast to our model where knowledge is treated as a public good (non-excludable and non-rivalrous). Together, these considerations suggest that depending on the nature of knowledge (e.g. public vs. common-pool resource), the traits or behaviours that underlie its acquisition may exhibit different levels of variation as they may experience different strengths of negative frequency-dependent selection.

Variation in traits related to knowledge acquisition has been reported across several taxa, e.g. birds [68–72], reptiles [73], fish [74], insects [75], and mammals [76] including humans [77]. In zebra finches, for instance, behavioral assays have revealed that females that forage less actively when in isolation tend to rely more heavily on social information for feeding preference when exposed to an exemplar individual, suggesting a negative association between individual and social learning [72]. Similarly in common marmosets, individuals that are better at using social information in one foraging task tend to be worse at finding optimal solutions individually at another task [76]. Such variation in social and individual learning could be due to genetic differences as in our model, and/or from a form of plasticity in learning (e.g., where learning behaviour depends on past experiences under unpredictable conditions [9]). Distinguishing between these two sources of variation would require estimating the heritability of social and individual learning, which remains a challenging task. Nevertheless, multiple studies on both humans and other animals point to genetic contributions to learning and associated cognitive skills (for review [78]).

Our results specifically point to a potential association between teaching and individual learning behaviours, which to the best of our knowledge has not yet been described in natural populations. Apart from humans, this idea could potentially be tested in Meerkats (*Suricata suricatta*) as variability in both teaching and individual learning behaviours has been documented in this species. For example, when adult meerkats present live prey to pups, 87.5% remain with the pup to monitor its handling of the prey, while 12.5% do not [45]. Additionally, variation has been observed in the time spent manipulating an apparatus during a food extraction task and the likelihood of successfully obtaining the food [79]. To test the potential association between teaching and individual learning, one could for instance quantify whether individuals that exhibit more teaching behaviour (e.g., remaining with pups to monitor prey handling) are less successful at solving individual learning tasks, such as a food extraction task.

More fundamentally, our results reinforce the general notion that kin selection plays a key role in the evolution of behaviours and structures related to the acquisition and transmission of knowledge and skills [10, 12, 26–31, 49–53]. In our model, kin selection effects are driven by vertical transmission of knowledge from parent to offspring, in line with the observation that the majority of documented cases of teaching in animals occur from parent, typically mothers, to offspring [35–37, 40, 43, 80]. Cases of oblique or horizontal teaching outside of humans have so far been limited to societies of cooperative breeders [81] and eusocial insects [32] both of which show high levels of relatedness. Teaching among non-relatives may evolve through mechanisms of reciprocity such that teachers obtain direct fitness benefits to offset teaching costs [53, 82]. An ethnographic study of Fijian hunter-gatherers revealed that although teaching occurs more frequently towards relatives in these societies, it is also prevalent among non-kin, especially to transmit skills deemed important for the success of village life [80]. This suggests that teaching may provide direct survival or reproductive benefits to teachers in that case, e.g. in the form of prestige deference exchange [83].

Our results are based on many assumptions, some of which merit further discussion. For simplicity, we assumed that individuals are haploid and reproduce asexually. Diploidy and sexual reproduction is expected to affect our results in two ways. First, meiotic recombination would break the positive association among traits that is favoured by selection, e.g. among teaching and individual learning here. However, if the traits’ genetic architecture can evolve (e.g., through recombination modifiers or pleiotropic loci), then synergistic fitness effects between traits should favour an architecture that allows these associations to be heritable [84]. Second, by reducing parent-offspring relatedness by one half, diploidy and sexual reproduction would favour correspondingly lower levels of teaching than in our model. But this effect would be quantitative rather than qualitative [27] so that our main results would remain unchanged. Another assumption we made is that individuals can only perform one type of learning at a time. While this is undoubtedly a simplification of reality—where individuals likely acquire knowledge and skills from multiple sources throughout their lives—it captures the general trend that social learning is predominant early in life, with individual learning becoming more prominent as individuals age [85–88]. In humans, there is also a clear shift in the focus of social learning over time, from predominantly vertical learning during childhood to increased reliance on oblique learning in later life stages [89, 90]. Our assumptions on learning captures such features in a stylised way to make the model tractable and we discuss further modelling assumptions in Appendix E.

In conclusion, our study highlights the evolutionary interplay between learning schedules and parental teaching as potentially relevant in the origins of traits related to the acquisition, transmission, and accumulation of knowledge. This interplay enhances innovation and inter-generational transfer of information, as well as contributes to variation in knowledge within populations, manifesting as an association between teaching and innovating. More broadly, our model underscores the role of parent-offspring interactions in driving the evolution of behaviours and structures that optimise information acquisition and transmission among individuals. This is likely to be relevant not only for the adaptation of many animal species, but may also lay the foundations for the unique and extensive cumulative knowledge of humans, as the abilities for knowledge transmission and communication that evolved in the inner circle of parent-offspring interactions may later be expanded to the outer circles of non-familial interactions through norms and institutions, which are so characteristic of modern human societies.

## Methods

As mentioned in section 2.3, our analyses are based on a quantitative genetics approach to the joint evolution of multiple traits under rare mutations with small trait effects, such that the trait distribution in the population is approximately multi-variate Gaussian with mean vector ***x*** and variance-covariance matrix ***G***. This allows us to ascertain the dynamics of ***x*** and ***G*** from the lineage fitness (or invasion fitness or geometric growth ratio) of a rare type ***x***_*•*_ *=* (*v*_*•*_, *o*_*•*_, *τ*_*•*_) in a population which can be considered to be essentially monomorphic for mean trait values ***x*** *=* (*v, o, τ*) [56]. Lineage fitness is equal to the expected number of successful offspring that are produced by an individual randomly sampled from its lineage. We characterise lineage fitness for our model and then specify how it can be used to track the evolutionary dynamics below.

### Lineage fitness

Consider a population expressing ***x*** *=* (*v, o, τ*). The dynamics of individual knowledge and total knowledge in the population settle on the equilibrium 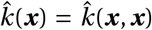 (given by the solution of eq. 5 with ***x***_*•*_ *=* ***x***) and 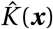 (given by eq. 6), respectively. Meanwhile, demographic dynamics settle for the equilibrium 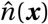, which is such that each individual in the population exactly replaces itself. This is found by solving 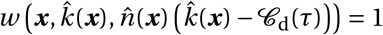 (eq. 4) for the equilibrium number of adults 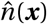, where

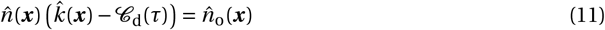

is the number of offspring that enter density-dependent competition according to our assumptions detailed in section 2.2. This yields

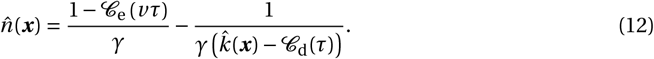

Consider now a lineage of individuals expressing ***x***_*•*_ *=* (*v*_*•*_, *o*_*•*_, *τ*_*•*_) in this population, which is assumed to be large. Individual knowledge in this lineage converges to the equilibrium 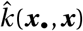 (solution of eq. 5) and the expected number of successful offspring that a lineage member produces at this equilibrium is given by 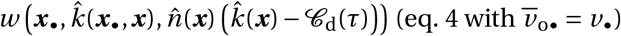 (eq. 4 with *v* _o*•*_ *= v*_*•*_). Plugging eq. (12) into this then yields the lineage fitness of type ***x***_*•*_

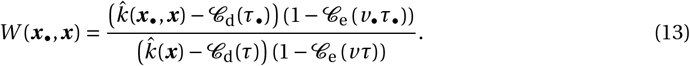

This determines whether the lineage grows (when *W* (***x***_*•*_, ***x***) *>* 1) or goes bust (when *W* (***x***_*•*_, ***x***) ≤ 1) in a population at ***x***.

Although it may not be immediately obvious from eq. (13), lineage fitness takes into account all the relevant kin selection effects brought about by vertical knowledge transmission. It does so via its dependency on equilibrium individual knowledge 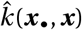. Indeed, from the point of view of a focal individual of that lineage (i.e. a randomly sampled member of this lineage), the individual knowledge 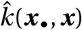 it possesses depends on the traits expressed by all its ancestors connected by vertical learning ([27] for more details).

### Dynamics of the mean trait vector *x*

For a given variance-covariance matrix ***G***, the change Δ***x*** *=* ***x***′ − ***x*** in the mean trait vector over one generation is given by

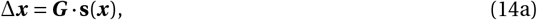

where **s**(***x***) *=* (*s*_*v*_ (***x***), *s*_*o*_(***x***), *s*_*τ*_(***x***)) is the selection gradient vector, pointing in the direction favoured by selection in trait space (here {(*v, o, τ*) : 0 ≤ *v* ≤ 1, 0 ≤ *o* ≤ 1, 0 ≤ *v +o* ≤ 1, *τ* ≥ 0}). The selection gradient is given by

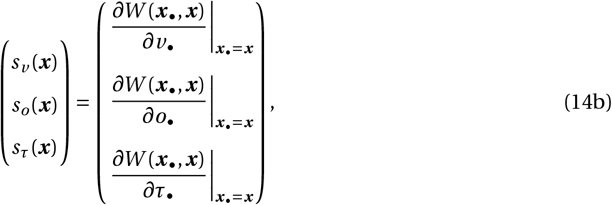

so that each entry determines whether selection favours an increase or decrease in the corresponding trait. As seen in eq. (14a), the response of trait means to selection over one generation is constrained by available variation captured by the ***G*** matrix [91]. When mutations are rare with small effects on trait values, changes in ***G*** occur more slowly than changes in ***x*** so that eq. (14a) can be considered with ***G*** constant [56, 92–95]. According to this eq. (14a), ***x*** eventually converges towards a convergence stable trait vector ***x***^*^ that either sits on the boundary of the trait space, or is in the interior of the trait space such that there is no more directional selection, i.e. such that **s**(***x***^*^) *=* (0, 0, 0).

### Dynamics of the trait variance-covariance matrix *G*

If ***x***^*^ is in the interior of the trait space, then the so-called Hessian matrix

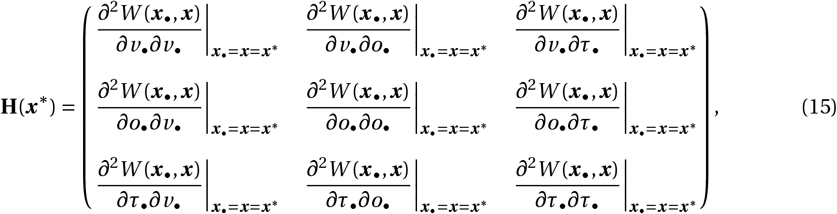

determines selection on the variance-covariance matrix ***G*** ([56, 93]; for details on the dynamics of ***G***, see eq. B.6 in Appendix). The dynamics of the variance-covariance matrix can have two outcomes depending on the matrix **H**(***x***^*^).

When the leading eigenvalue of **H**(***x***^*^) is negative, selection is stabilising around ***x***^*^, such that the trait distribution in the population converges under mutation-selection to be approximately Gaussian with mean ***x***^*^ and variance-covariance matrix 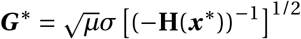, where *µ* is the probability of a mutation, and *σ*^2^ is the variance in the effect of a mutation on trait value, here assumed to be identically and independently distributed for each trait (the operation **X**^1/2^ denotes the square root of a matrix **X**, defined as the unique positive semidefinite and symmetric matrix that satisfies **X**^1/2^**X**^1/2^ *=* **X**, e.g. eq. (11) in [56]).

When the leading eigenvalue of **H**(***x***^*^) is positive, selection is disruptive and favours an increase in trait variance in trait space along the eigenvector associated with the leading eigenvalue of **H**(***x***^*^) [96, 97]. This may lead to evolutionary branching whereby the phenotypic distribution goes from being unimodal to bimodal so that two highly differentiated morphs coexist in the population.

### Overview of analyses

The above provides a recipe to investigate the joint evolutionary dynamics of teaching and learning schedules, and its effect on knowledge and demography. Our analyses following this recipe are detailed in Appendix C. Briefly, we first substitute eq. (13) (together with the solution of eq. 5) into eq. (14) to determine the equilibrium mean vector ***x***^*^ favoured under directional selection. If ***x***^*^ is in the interior of the trait space, we then substitute eq. (13) into eq. (15) to determine whether selection is stabilising (in which case we characterise the equilibrium variance-covariance matrix ***G***^*^), or disruptive (in which case we use individual-based simulations to determine whether evolutionary branching occurs). In addition, we use ***x***^*^ to characterize individual and total knowledge in the population at an evolutionary equilibrium from 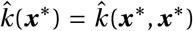 (solution of eq. 5) and 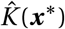 (eq. 6). This eq. (6) provides an excellent approximation for 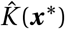 provided individuals perform some social learning (i.e. *v +o >* 0). To ensure this is always the case, we assume that in the absence of teaching (i.e. *τ =* 0 is fixed in the population), selection favours at least some social learning. This requires that either

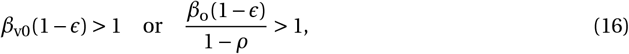

holds (Appendix C.2 for derivation and interpretation).

## Code availability

All the codes used in this study are accessible at github.com/Ludovic-Maisonneuve/coevo_learning_teaching.

## AI declaration

LM has used AI tools to spot grammar and spelling mistakes.

## Acknowledgments

LM would like to thank Iris Prigent, Arthur Weyna, Cédric Perret and Thomas Lesaffre for the useful discussions.

## Appendix to

### A Model of knowledge acquisition

In this Appendix A, we detail the assumptions underlying knowledge acquisition and its dynamics in our model.

#### A.1 Social learning as a foraging process

##### A.1.1 Searching and handling information

We conceptualise acquiring adaptive information as a foraging process that requires: (i) searching for/finding new information (e.g. seeking an exemplar individual, unknown piece of information); and (ii) handling new information (e.g. memorising, practising to make it work). Once a unit of information is handled, it becomes a unit of the knowledge of the learner. Accordingly, we can use classical foraging theory [1] and write the rate at which an individual learns a unit of knowledge per unit time via social learning generically as

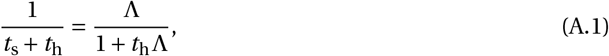

where *t*_s_ is the average time needed to search for a new unit of information, *t*_h_ is the average time needed to handle that unit of information; and Λ *=* 1/*t*_s_ is then the rate at which an individual encounters a new unit of information (i.e. a unit it does not already know). The encounter rate Λ is assumed to be proportional to the amount of available knowledge in the population that the foraging individual does not already have.

##### A.1.2 Encounter rate and knowledge dynamics

We next characterise Λ for the vertical and oblique learning stages and derive eq. (1) of the main text. To that aim, we focus on a particular generation *t* and let *k*_*•t*_ (*a*) ∈ [0, ∞) denote the knowledge of a focal offspring from that generation at age *a* ∈ [0, 1], and where this focal individual comes from a lineage of individuals all expressing traits ***x***_*•*_ *=* (*v*_*•*_, *o*_*•*_, *τ*_*•*_). We assume that individuals start their lives with zero knowledge so that *k*_*•*_(0) *=* 0.

###### Vertical learning

Following our assumption (section 2.1), the focal offspring first learns vertically from its parent for a period of length *v*_*•*_. During this phase, the rate Λ_v_ at which the focal encounters novel up-to-date information is

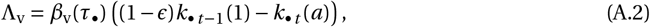

where recall *ϵ* is the rate at which social knowledge becomes obsolete and *k*_*•t*−1_(1) is the knowledge of the focal’s parent (which corresponds to the parent’s knowledge at the end of its learning phase, i.e. at age *a =* 1). Hence, the quantity (1 − *ϵ*)*k*_*•t*−1_(1) − *k*_*•t*_ (*a*) is the amount of up-to-date parental knowledge that the offspring does not already have at age *a*. The vertical transmission rate *β*_v_(*τ*_*•*_) is given by eq. (2).

###### Oblique learning

Next, the focal offspring learns obliquely from all adults in the population for a period of time *o*_*•*_. During this phase, the rate Λ_o_ at which it encounters novel up-to-date information is

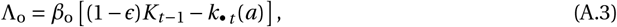

where *K*_*t*−1_ is the total amount of information that is known by all the adults from the previous generation and that the focal offspring has access to through social learning. We refer to this as the “total knowledge” for short and model its dynamics in Appendix A.2.

###### Individual learning

After learning socially, individuals produce new knowledge at rate *α*, and the focal offspring does so for a period of length 1 − *v*_*•*_ −*o*_*•*_.

Substituting eq. (A.2) and eq. (A.3) into eq. (A.1), we obtain the rate of learning, d*k*_*•t*_ (*a*)/d*a =* 1/(*t*_s_ *+ t*_h_), of the focal individual during vertical and oblique learning, respectively. Putting it all together with individual learning, we get that the rate of learning of the focal individual over its lifetime follows the ordinary differential equation:

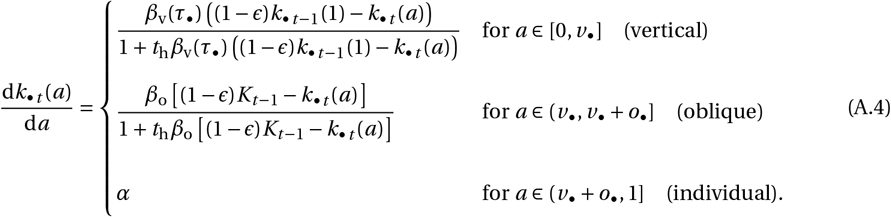

In general, eq. (A.4) cannot be solved explicitly for *k*_*•t*_ (*a*). For the bulk of our analysis, we therefore assume that the handling time is negligible. We explore the impact of handling time on evolutionary dynamics in Appendix D, where we show that handling time has limited effects unless it is so large that it disfavours the evolution of social learning altogether. With *t*_h_ *=* 0, eq. (A.4) reduces to eq. (1) of the main text. Note that with *t*_h_ *=* 0, the net rate of social learning is proportional to the amount of available knowledge an individual does not already have. Previous models making this assumption (e.g., [2–7]) can therefore be seen as model of information foraging with negligible handling time. By contrast, when *t*_h_ is large compared to *t*_s_, the learning rate becomes constant (as search time is no longer the limiting factor). In this case, the oblique learning part of eq. (A.4) becomes qualitatively equivalent to that found in [8].

#### A.2 Total knowledge dynamics

We now specify the dynamics of the total knowledge in the population and derive eq. (3) of the main text. For our purpose, it is sufficient to do so under the assumption that the population is effectively monomorphic for ***x*** *=* (*v, o, τ*). This means that all offspring have the same learning schedule and therefore accumulate knowledge according to eq. (A.4) with ***x***_*•*_ *=* ***x***. Let 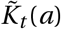 be the total knowledge accumulated by the offspring of age *a* in generation *t* that successfully establish as adults, of which there are *n*_*t*_. The total knowledge among the adults of generation *t*, *K*_*t*_, is then given by 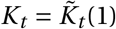.

Using this notation, we model 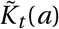 such that it satisfies the ordinary differential equation

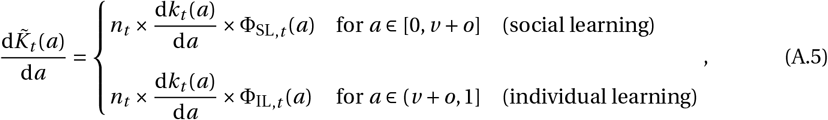

with initial condition 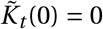, where d*k*_*t*_ (*a*)/d*a* is given by eq. (A.4) with ***x***_*•*_ *=* ***x***, and 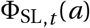 and 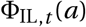, are the proportion of the knowledge learned by an individual at age *a* that increase total knowledge during social and individual learning, respectively (so that they are both between zero and one). These terms account for potential redundancy in learning among individuals. For instance, if during social learning an individual acquires a unit of information that is already present in the population because others previously acquired them socially, this does not increase the total knowledge, and 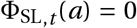. Similarly, if during individual learning the *n*_*t*_ individuals simultaneously all produce the same unit of knowledge, this unit should only be counted once in the total knowledge, and 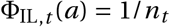. We next specify how we derive the expressions of 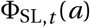 and 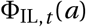 from the assumption of the model of knowledge acquisition.

During social learning, the total knowledge grows when an offspring learns a unit of knowledge which has not yet been passed onto any of the *n*_*t*_ offspring. 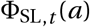 is then given by the proportion of knowledge available for an offspring by social learning that has not been transmitted yet to any of the *n*_*t*_ offspring, i.e. by

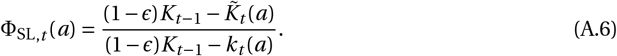

The numerator is the up-to-date total knowledge of the previous generation that has not been transmitted to any of the *n*_*t*_ offspring, and the denominator is the up-to-date total knowledge of the previous generation that an offspring has yet to learn. The amount of total knowledge accumulated via social learning at generation *t* is then given by 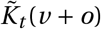 as the solution of eq. (A.5) with eq. (A.6) substituted into it. Unfortunately, we cannot obtain such a solution analytically. However, note that if *n*_*t*_ is large, then the right hand side of the first line of eq. (A.5) (with eq. A.6) is large. As a result, 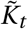should quickly converge to (1 −*ϵ*)*K*_*t*−1_, which is amount of total knowledge that is available from social learning. Provided the population is large, we can therefore use the approximation

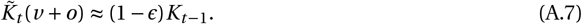

Fig. A.1 shows that this is an excellent approximation to the exact dynamics with *n*_*t*_ as small as 100. After social learning, offspring produce knowledge independently via individual learning, which in-creases total knowledge. We assume that during individual learning, an individual produces units of information that are not already known among the other individuals of the same generation, but it may produce the same unit of information as another individual also performing individual learning. With *ρ* as the probability that two individuals produce different pieces of knowledge when learning individually simultaneously (recall section 2.1), we then have

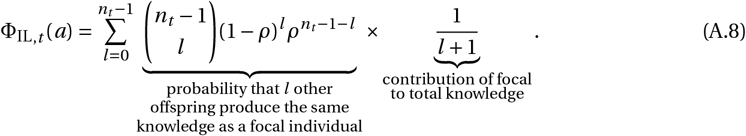

Applying the binomial theorem and simplifying we get

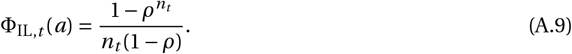

which allows us to compute the amount of total knowledge accumulated at the end of the learning phase in generation *t*. Indeed, the accumulated knowledge at the end of the learning phase using eq. (A.5) is given by

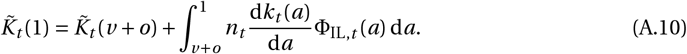

Substituting eq. (A.9) and d*k*_*t*_ (*a*)/d*a = α* into the above, we obtain

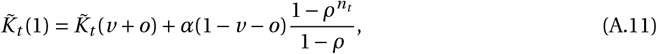

where the second term can be written as

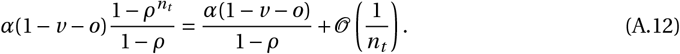

Using eq. (A.7), eq. (A.11) and eq. (A.12) and recalling that 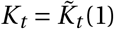, we get that the total knowledge at generation *t* can be approximated by

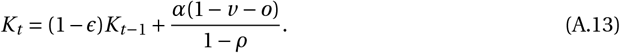

Substituting this into Δ*K = K*_*t*_ −*K*_*t*−1_ then obtains eq. (3) of the main text, as required.

#### A.3 Equilibrium

Finally, we determine the equilibrium of the knowledge dynamics under vanishing handling time and derive eq. (5) of the main text. Solving eq. (A.4) with *t*_h_ *=* 0 and 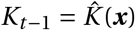 allows us to obtain the knowledge that a focal offspring bears at the age of reproduction in generation *t* as

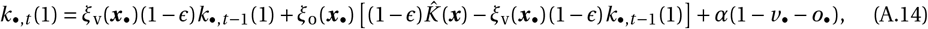

Where

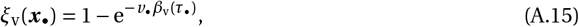

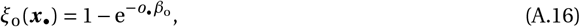

(see below main text eqs. 5 and 6 for an interpretation of *ξ*_v_(***x***_*•*_) and *ξ*_o_(***x***_*•*_)). From eq. (A.14), it is then straightforward to obtain that the equilibrium 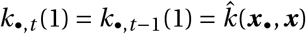 is

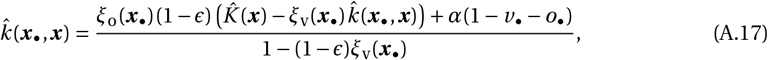

which is eq. (5) of the main text, as required, and where

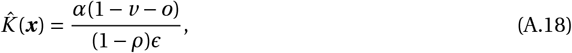

is found by solving eq. (A.13) at 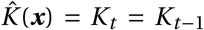 (and is eq. (6) of the main text). Substituting eq. (A.18) into eq. (A.17) and re-arrangements then yield

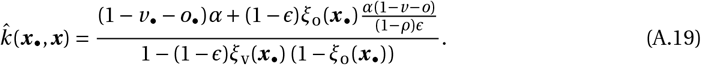

### B. Mathematical evolutionary approach

Here, we describe in greater detail our evolutionary approach that is outlined in the Methods section. Our main assumption is that the three traits *v*, *o* and *τ* evolve via rare small effect mutations such that the multi-trait distribution in the population at any generation *t* is approximately multi-variate

**Figure A.1:**
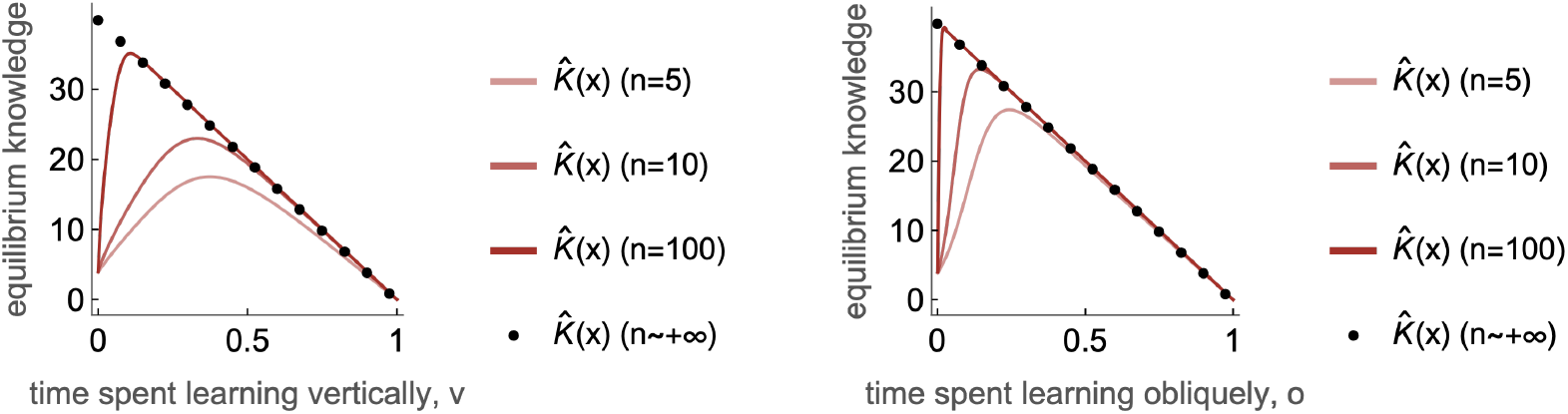
Equilibrium total knowledge. 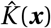. Red curves are obtained by computing numerically 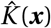 using eq. (A.5), (A.6) and (A.9) for different constant population sizes *n* (see figure for values). Black dots are obtained using eq. (A.18), which assumes a large population. Parameters: *v =* 0, *o =* 0, *τ =* 0.1, *β*_v0_ *=* 7, *β*_τ_ *=* 10, *β*_o_ *=* 4, *α =* 2, *ϵ =* 0.1, *ρ =* 0.5)

Gaussian 𝒩 (***x*** _*t*_, ***G*** _*t*_) where

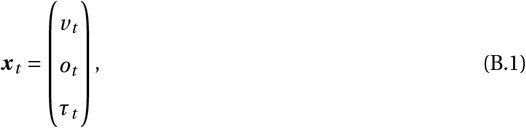

is the vector of trait means, and

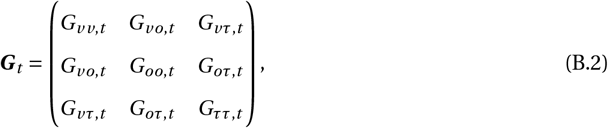

is the symmetric variance/covariance matrix, with *G*_*yz*,*t*_ *= G*_*zy*,*t*_ denotes the covariance among traits *y* and *z* (both in {*v, o, τ*}) at generation *t*. The dynamics of the phenotypic distribution is determined by the dynamics of the vector ***x*** _*t*_ and of the matrix ***G*** _*t*_. Where mutations have weak effects such that (co)variances are small, these dynamics can be studied separately owing to a timescale decomposition between changes in the means (which are fast) and in the (co)variances (which are slow, e.g. [9–13]). We detail these changes in the next two sections.

#### B.1 Directional selection

First, the population evolves under directional selection whereby the vector of mean trait values ***x*** _*t*_ changes according to

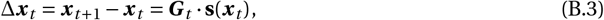

where ***G*** _*t*_ is the variance/covariance matrix and

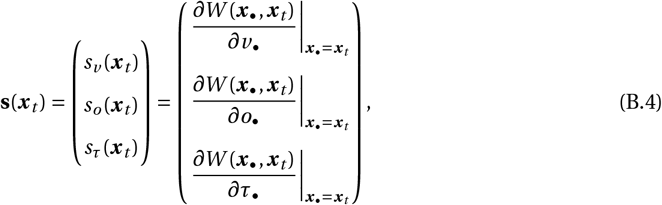

is the selection gradient, which is a vector pointing in the direction favoured by selection in phenotypic space, here {(*v, o, τ*) : 0 ≤ *v* ≤ 1, 0 ≤ *o* ≤ 1, 0 ≤ *v + o* ≤ 1, *τ* ≥ 0} [13–16]. Specifically, the selection gradient consists of the marginal effects of each trait on lineage fitness, *W* (***x***_*•*_, ***x*** _*t*_), which is given by eq. (13) with eq. (A.19).

When phenotypic variance is small, the vector of mean trait values evolves faster than the variance/covariance matrix. This entails that the mean trait values may converge under directional selection to an equilibrium before any significant changes occur in the variance/covariance matrix. Therefore, the variance/covariance matrix can be considered constant (***G*** _*t*_ *=* ***G***) during this initial phase of evolutionary dynamics. Assuming that each trait is encoded by a separate locus and all loci mutate in a similar way, it is reasonable to put ***G*** *= δ****I*** here with *δ >* 0 a small constant and ***I*** the identity matrix.

The evolutionary dynamics under directional selection, given by eq. (B.3) with ***G*** _*t*_ *=* ***G***, may eventually converge to an equilibrium, ***x***^*^, which in this case is referred to as a convergence stable strategy or convergence stable trait vector [17–22]. From eq. (B.3) with ***G*** _*t*_ *=* ***G*** and using the fact ***G*** is a variance-covariance matrix and thus a positive-definite matrix, it is necessary that

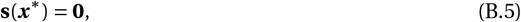

for ***x***^*^ in the interior of phenotypic space to be convergence stable.

#### B.2 Stabilising, disruptive and correlational selection

Once average trait values have converged to a convergence stable trait vector ***x***^*^, the variance-covariance matrix ***G*** _*t*_ evolves according to

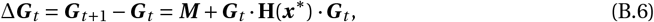

Where

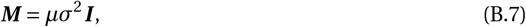

is a constant positive-definite matrix that reflects the input of uncorrelated mutations (which occur with probability *µ* and whose effect on a trait value has a variance *σ*^2^), and

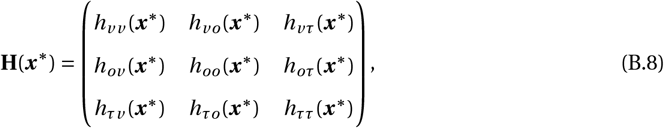

is the so-called Hessian matrix whose entries are defined as

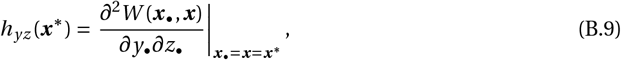

for each pair of traits (*y, z*) with *y, z* ∈ {*v, o, τ*} [13]. The diagonal entries of the Hessian matrix indicate whether selection on each individual trait *y* is stabilising (*h*_*y y*_ (***x***^*^) *<* 0) or disruptive (*h*_*y y*_ (***x***^*^) *>* 0), resulting respectively in a reduction or an increase in the trait’s variance when the trait evolves in isolation from the others [23]. The off-diagonal entries, meanwhile, indicate whether selection favours a positive (*h*_*yz*_ (***x***^*^) *>* 0) or a negative (*h*_*yz*_ (***x***^*^) *<* 0) correlation among two traits *y* and *z* (with *y≠ z*). Accordingly, these off-diagonal entries have been coined as the coefficients of correlational selection [23].

The dynamics given by eq. (B.6) can have two outcomes depending on the leading eigenvalue of **H**(***x***^*^).

When the leading eigenvalue of **H**(***x***^*^) is negative, selection is stabilising, keeping traits’ (co)variances small so that the phenotypic distribution remains unimodal around the equilibrium [22]. The variance/covariance matrix converges towards an equilibrium ***G***^*^ determined by the joint action of mutation and selection; specifically,

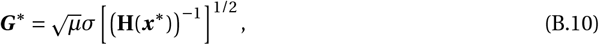

where the operation **X**^1/2^ denotes the square root of **X** such that all the eigenvalues of **X**^1/2^ are positive (see eq. (11) in [13]).

When the leading eigenvalue of **H**(***x***^*^) is positive, selection becomes disruptive, increasing trait variance in trait space along the eigenvector associated with the dominant eigenvalue of **H**(***x***^*^). This may lead to evolutionary branching, whereby the trait distribution in the population goes from being unimodal to bimodal [22].

### C Evolutionary analysis

#### C.1 Selection gradient

Using Equations (14b) (or B.4), (13) and (A.19) we obtain the selection gradient for *v*, *o* and *τ*:

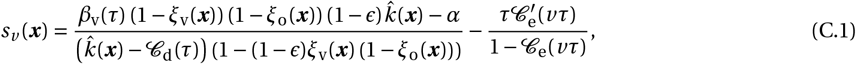

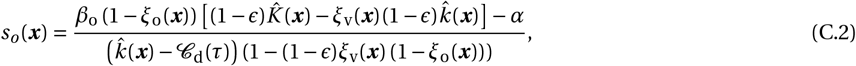

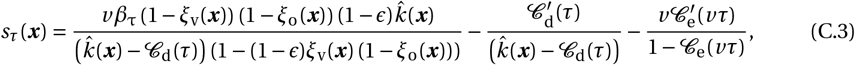

where 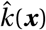 is given by eq. (A.19) with 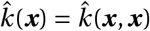, and 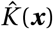 is given by eq. (A.18).

#### C.2 Emergence of social learning

Here, we derive eq. (16) of the main text, which specify the conditions for the emergence of vertical or oblique learning in the absence of teaching. The conditions for vertical or oblique learning to emerge are respectively given by

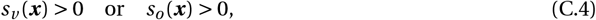

with ***x*** *=* (0, 0, 0). Using the expression of the selection gradient for *v* and *o* given in eq. (C.1)and eq. (C.2) with *v = o = τ =* 0, condition (C.4) becomes

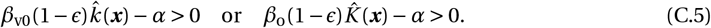

To express the condition (C.5) in terms of the model parameters, we need to compute and substitute the expressions for 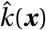 and 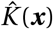. From eq. (A.19) with ***x***_*•*_ *=* ***x*** *=* (0, 0, 0), we obtain that

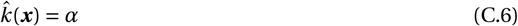

When *v = o =* 0, there is no transmission of knowledge across generations. Consequently, the total knowledge in each generation is equal to the knowledge produced within a single generation which is given in eq. (A.12). Using eq. (A.12) with *v = o =* 0, we obtain that

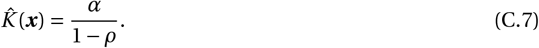

Plugging eqs. (C.6) and (C.7) into eq. (C.5), we then have

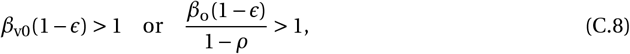

which is eq. (16) of the main text, as required.

#### C.3 Evolution in the absence of teaching

Throughout this section, we fix *τ =* 0 so that there is no teaching in the population.

##### C.3.1 Evolutionary equilibrium where individuals perform all three types of learning

Here, we derive eq. (7) of the main text. At the evolutionary equilibrium 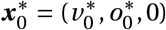, individuals perform all three types of learning if 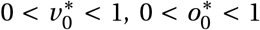, and 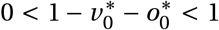 all hold. For individuals to perform all three types of learning at 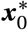 it is necessary that 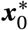 satisfies

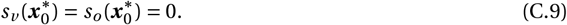

By substituting eq. (C.1) and eq. (C.2) with 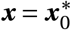 into eq. (C.9), it becomes

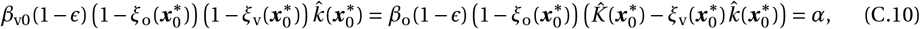

where 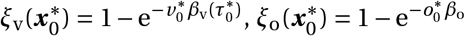 and where the expression of 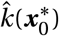 and 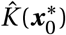 can be obtained using 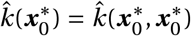 and by substituting 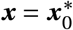 into eqs. (A.19) and (A.18), respectively. eq. (C.10) is eq. (7) of the main text, as required.

##### C.3.2 Marginal benefits of vertical, oblique, and individual learning

We demonstrate that eq. (7) is equivalent to the condition that for a focal individual, the marginal effects of increasing its time learning vertically, obliquely and individually on its knowledge at reproduction are all equal. To see this, let’s consider the knowledge *k*(1) at the end of the learning period of an individual who extends the duration of its vertical, oblique, and individual learning phases by *δ*_*v*_, *δ*_*o*_, and *δ*_*i*_, respectively in comparison to a focal individual in a population expressing 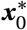. By defining *k*(1) *= k*_*•*_(1 *+ δ*_*v*_ *+ δ*_*o*_ *+ δ*_*i*_) using eq. (1) with 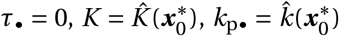, *v*_*•*_ *= v*^*^ *+ δ*_*v*_, *o*_*•*_ *= o*^*^ *+ δ*_*o*_ and by letting the phase of individual learning last until *a =* 1 *+δ*_*v*_ *+δ*_*o*_ *+δ*_*i*_, we obtain

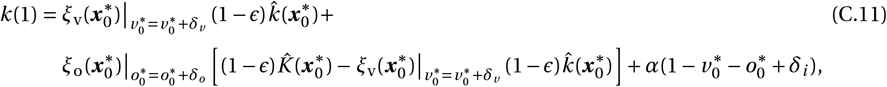

where 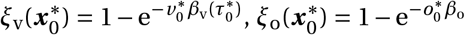 and where the expression of 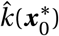 and 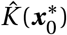 can be obtained using 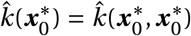, and by substituting 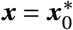 into eqs. (A.19) and (A.18), respectively. With this notation, eq. (7) is equivalent to

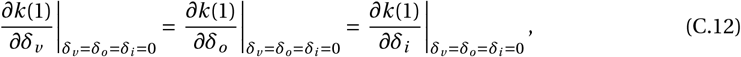

which shows that each term in equation (7) represents the marginal effect of an increase in the time spent in each learning phase, while the duration in the other phases remains unchanged.

##### C.3.3 Necessary conditions for the evolution of all three types of learning

We derive here eq. (8) of the main text, which gives the necessary condition for the existence of an evolutionary equilibrium where individuals perform all three types of learning. To that end, we investigate when eq. (C.10) has a solution such that 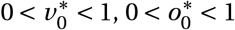, and 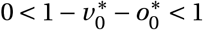.

First, note from eqs. (C.1) and (C.2) (and eq. (A.19) with 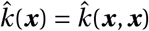 and eq. (A.18)) with *τ =* 0 that

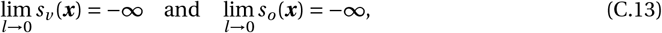

where *l =* 1 − *v* − *o* is the time spent on individual learning. eq. (C.13) shows that selection always favours some time allocated to individual learning, i.e., that 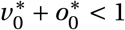. We therefore only need to investigate whether eq. (C.10) can have a solution such that 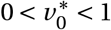 and 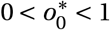.

Now, the conditions 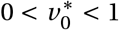 and 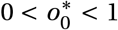 hold if, and only if, 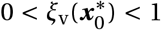 and 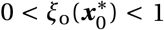 (where 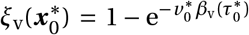 and 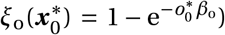). Let us then consider the first equality in eq. (C.10), which can be re-arranged as

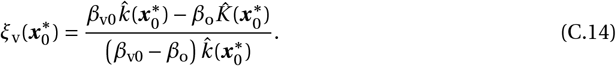

Plugging eq. (C.14) into the condition 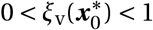, we obtain that it is necessary that

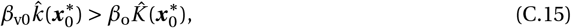

for 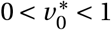1 to hold, which is the first of condition (8) of the main text.

Finally, we consider the second equality in eq. (C.10). This can be re-arranged to read as

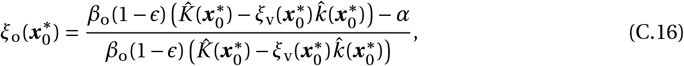

which plugged into the condition 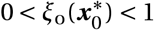 reveals that it is necessary that

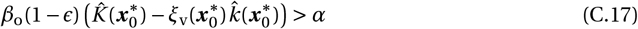

for 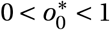 to hold. However, substituting for 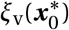 from eq. (C.14), this can be re-written as,

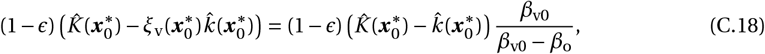

which is the second of condition (8) of the main text, as required.

##### C.3.4 Numerical analysis of the dynamics of the mean trait vector

To determine numerically the evolution of the mean trait vector ***x***, we used eq. (B.3) to iterate

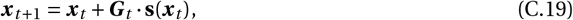

for *t* ≥ 0 starting with ***x***_0_ *=* (0, 0, 0), where the selection gradient is given by eqs. (C.1)–(C.3) (using eq. A.19 and eq. A.18), and the variance-covariance matrix is fixed at

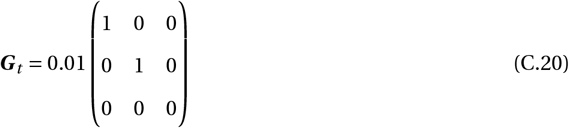

for all *t*, to ensure that *τ =* 0 always. When necessary, we truncate ***x*** _*t*_ to remain in the trait space {(*v, o, τ*) : 0 ≤ *v* ≤ 1, 0 ≤ *o* ≤ 1, 0 ≤ *v + o* ≤ 1, *τ* ≥ 0}. We iterate eq. (C.19) until ∥***x*** _*t+*1_ − ***x*** _*t*_ ∥ *<* 10^−7^. This method was used to make Figs. 2, 3, S1, S3, and S4.

#### C.4 Coevolution of learning schedules and teaching

##### C.4.1 Condition for the emergence of teaching

Here we detail how we get eq. (9) of the main text, which gives the condition for the emergence of teaching. We assume that the population has reached the evolutionary equilibrium 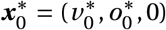. For teaching to evolve within this population, it is necessary that

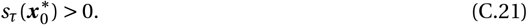

By substituting eq. (C.3) with 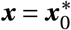 into eq. (C.21), we obtain the condition

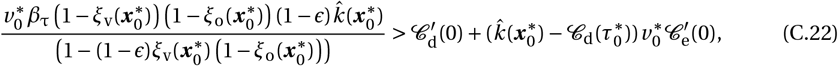

where 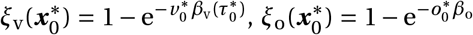 and where the expression of 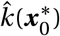 can be obtained using eq. (A.19) with 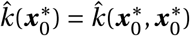. We can then use the fact that

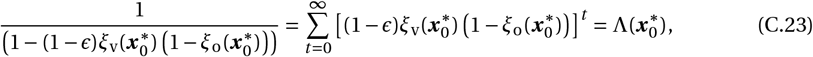

to obtain condition (9) of the main text, as required.

##### C.4.2 Effect of teaching on the inter-generational vertical transmission rate

Here we show that 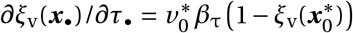, which we use in the main text (section 3.2). This is straightforward as from 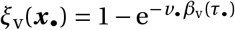 and eq. (2), we have

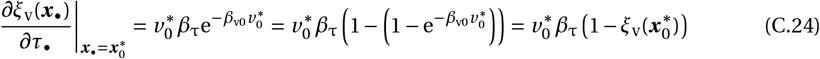

where we used 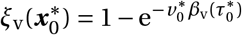 for the third equality.

##### C.4.3 Numerical analysis when learning schedules and teaching coevolve

To determine the evolution of the mean vector of traits ***x*** when all three traits coevolve, we use the same procedure exposed in section C.3.4, except that to allow teaching to evolve, the variance-covariance matrix now is

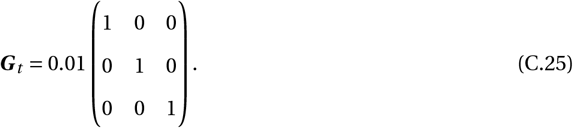

This method was used in the making of Figs. 3, 4, S3, S4 and S5.

#### C.5 Individual-based simulations

##### C.5.1 Algorithm

Our individual-based simulations track the evolution of the trait distribution, the distribution of individual knowledge, the total knowledge, and the population size at each generation for a fixed number of generations following the life cycle described in the main text. Each individual *i* at each generation is characterised by the vector of its three traits (*v*_*i*_, *o*_*i*_, *τ*_*i*_), as well as the teaching effort *τ*_p,*i*_ and the knowledge *k*_p,*i*_ of its parent. At the first generation, the values of *v*_*i*_, *o*_*i*_, *τ*_*i*_ and *τ*_p,*i*_ are initialised to *v* ^1^, *o*^1^, *τ*^1^ and *τ*^1^ for each individual *i* ∈ {1,…, *n*_1_}, where *n*_1_ *=* 1000 is the number of individuals generation 1. Additionally, *k*_p,*i*_ is set to 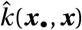 (from eq. A.19) with ***x***_*•*_ *=* ***x*** *=* (*v* ^1^, *o*^1^, *τ*^1^), and *K*_0_ in the first parental generation (generation 0) is given by 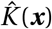 (from eq. A.18) with ***x*** *=* (*v* ^1^, *o*^1^, *τ*^1^).

At each generation *t* ≥ 1, the following occurs:

###### (i) Learning

We first determine the knowledge *k*_*i*_ (1) of each adult *i* at the age of reproducing, for *i* ∈ {1, …, *n*_*t*_} where *n*_*t*_ is the number of adult individuals at generation *t*. We do so by solving numerically the ordinary differential equation

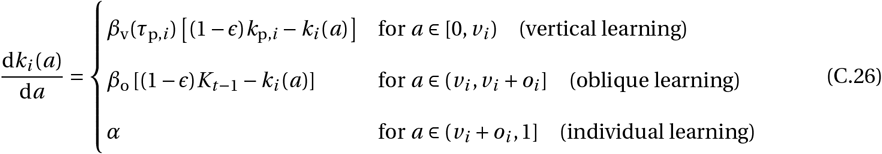

with *k*_*i*_ (0) *=* 0 for each *i* ∈ {1, …, *n*_*t*_} (eq. C.26 is eq. 1 with the notation introduced in the above paragraph).

###### (ii) Total knowledge

We calculate the total knowledge *K*_*t*_ among the adults of generation *t*, which is used in the next generation *t +* 1. We do this in two steps. First, we order individuals from 1 to *n*_*t*_, according to the amount of time each individual spends learning socially (and thus in descending order of the amount of time spent learning individually). To capture this ordering mathematically, let us define *g* as a function from the set {1, …, *n*_*t*_} to itself, such that the sequence 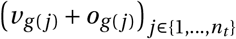 is increasing with *j*. Total knowledge *K*_*t*_ is then calculated as

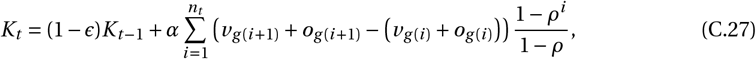

where we set 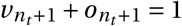. This eq. (C.27) is analogous to eq. (A.13), but unlike eq. (A.13), which assumes weak trait variance in the population, eq. (C.27) applies more generally including to the case where individuals have highly differentiated learning schedules (we derive eq. C.27 in section C.5.2 below).

###### (iii) Reproduction

The fecundity *f*_*i*_ of each individual *i* ∈ {1, …, *n*_*t*_} is calculated as *f*_*i*_ *= k*_*i*_ (1) −𝒞_d_(*τ*_*i*_) (first factor of eq. 4) with 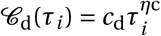. Each individual *i* then produces a number of offspring that is sampled from a Poisson distribution with mean equal to *f*_*i*_. In total, they produce a total number of offspring that we denote as *n*_o,*t*_.

###### (iv) Mutation

With probability 1 − *µ*, an offspring inherits the same traits as its parent. Otherwise (with probability *µ*), traits mutate; we model this by adding to each parental trait value an effect sampled from a normal distribution with mean 0 and variance *σ*^2^. If necessary, the resulting trait values are truncated to remain in {(*v, o, τ*) : 0 ≤ *v* ≤ 1, 0 ≤ *o* ≤ 1, 0 ≤ *v +o* ≤ 1, *τ* ≥ 0}. In all simulations, we set *µ =* 0.002 and *σ =* 0.01.

###### (v) Density-dependent competition

The competitiveness *c*_*i*_ of each offspring *i* in {1, …, *n*_o,*t*_} is calculated as *c*_*i*_ *=* (1 −𝒞_e_(*v*_*i*_ *τ*_p,*i*_))/(1 *+γn*_o,*t*_) (second factor of eq. 4), where recall *n*_o,*t*_ is the total number of offspring produced (from step (ii) above), 𝒞_e_(*v*_*i*_ *τ*_p,*i*_) *= c*_e_(*v*_*i*_ *τ*_p,*i*_)^*η*e^ and *τ*_p,*i*_ is the teaching effort of the parent of *i*. Each offspring *i* then survives till adulthood with probability *c*_*i*_ (and otherwise dies). This results in the number *n*_*t+*1_ of adults of the next generation (with *n*_*t+*1_ ≤ *n*_o,*t*_).

We repeat steps (i)-(v) for a fixed number of generations (see figure legends for parameter values).

##### C.5.2 Dynamics of total knowledge under arbitrary trait variance

Here, we derive eq. (C.27). Recall from Appendix A.2 that 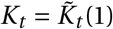 where 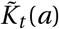 is the total knowledge accumulated by the offspring of age *a* in generation *t* that successfully establish as adults. This 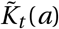 satisfies the ordinary differential equation

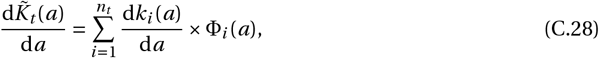

with initial condition 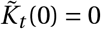, where *n*_*t*_ is the number offspring that will successfully establish and thus constitute the adults of the generation *t*, <u>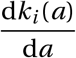</u> is the learning rate of individual *i* (given by eq. 1 with *v*_*•*_ *= v*_*i*_, *o*_*•*_ *= o*_*i*_, *τ*_p*•*_ *= τ*_p,*i*_, *k*_p*•*_ *= k*_p,*i*_ and *K = K*_*t*−1_), and Φ_*i*_ (*a*) is the proportion of the knowledge learnt by individual *i* at age *a* that contributes to total knowledge (same concept as in section A.2). From the definition that 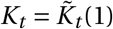, we thus have

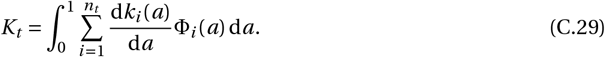

It is useful to distinguish between contributions from social and individual learning to total knowledge as

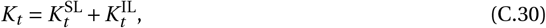

Where

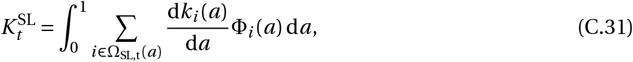

And

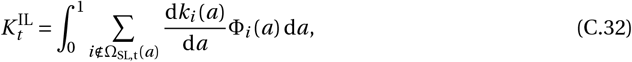

where Ω_SL,t_(*a*) is the set of individual among the *n*_*t*_ future adults that perform social learning (either vertical or oblique) at age *a* at generation *t*. Hence, the sum in eq. (C.31) is on all individuals performing social learning at age *a*, while the sum in eq. (C.32) is on all individuals performing individual learning at age *a*. Note that using the same argument as the one used to derive eq. (A.7), we can set

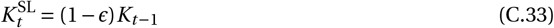

provided that the population is large enough.

Next, we decompose 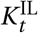 (eq. C.32) as

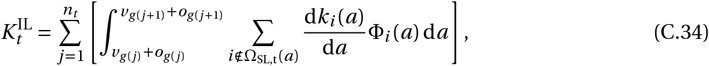

where recall from eq. (C.27) that *g* is a function such that 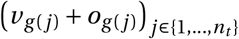 is increasing (setting 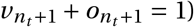). eq. (C.34) decomposes the integral in eq. (C.32) into the sum of *n*_*t*_ integrals such that during the time over which each integral is taken, the number of individuals performing individual learning remains constant. Specifically, provided *a* ∈ [*v*_*g*(*j*)_ *+ o*_*g*(*j*)_, *v*_*g*(*j+*1)_ *+ o*_*g*(*j+*1)_) in eq. (C.34), the number of individuals performing individual learning is *j*. Thus, using the same argument as the one used for eq. (A.9), we obtain that the share of knowledge Φ_*i*_ (*a*) learned by each individual *i* ∉ Ω_SL,t_(*a*) during individual learning at age *a* ∈ [*v*_*g*(*j*)_ *+ o*_*g*(*j*)_, *v*_*g*(*j+*1)_ *+o*_*g*(*j+*1)_] contributing to the total knowledge is

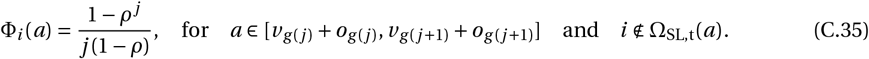

By injecting eq. (C.35) into eq. (C.34), and by using the fact that d*k*_*i*_ (*a*)/d*a = α* when individual *I* performs individual learning, we get

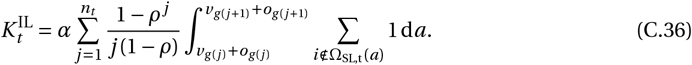

Now, since the number of individual performing individual learning is *j* during the time period *a* ∈ [*v*_*g* (*j*)_ *+o*_*g* (*j*)_, *v*_*g* (*j+*1)_ *+o*_*g* (*j+*1)_], we have

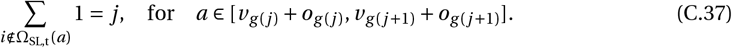

By plugging eq. (C.37) into eq. (C.36), we get

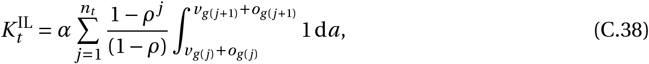

which simplifies to

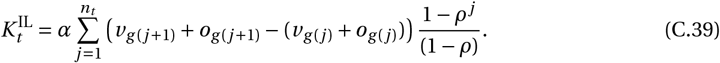

Finally, by substituting eqs. (C.33) and (C.39) into eq. (C.30), we obtain eq. (C.27), as required.

#### C.6 Densities of knowledge producers and scroungers

Here we show that after polymorphism emerges, the equilibrium density of knowledge producers tends to be very low, leading to their stochastic extinction. To study the dynamics of the number of producers and scroungers *n*_p_ and *n*_s_, we assume that after the population has reached the convergence stable trait vector ***x***^*^ and become polymorphic, the trait distribution is given by the sum of two Gaussian distributions, one for each morph and each of which shows small variance. With weak variance in each morph, evolutionary dynamics are slow relative to demographic changes [24, 25]. We can thus study the equilibrium numbers of producers *n*_p_ and scroungers *n*_s_ in the population assuming the mean trait values for each morph are fixed. Let us denote those by ***x***_p_ *=* (*v*_p_, *o*_p_, *τ*_p_) for the producer morph, and ***x***_s_ *=* (*v*_s_, *o*_s_, *τ*_s_) for the scrounger morph.

The population reaches demographic equilibrium when scroungers and producers both have the same fitness such that each individual exactly replaces itself, i.e. the equilibrium number of producers and scroungers 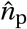 and 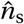 are such that

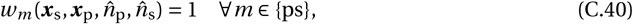

where *w*_*m*_(***x***_p_, ***x***_s_, *n*_p_, *n*_s_) is the expected number of offspring produced by an individual from morph *m* ∈ {p, s}, when there are *n*_p_ and *n*_s_ producers and scroungers in the population. Using eq. (4), this fitness is given by

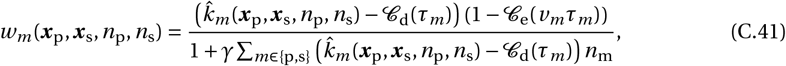

Where 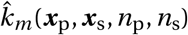 is the equilibrium knowledge in a lineage of individuals of morph *m* ∈ {p, s}. From eq. (A.19), this is

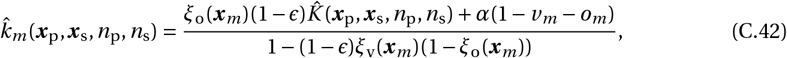

where 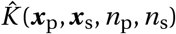 the equilibrium total knowledge in a dimorphic population.

To determine 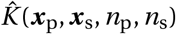, we can use eq. (C.27), which allows for an arbitrary distribution of traits in the population. For simplicity, we ignore variance within morphs so that there are two types of individuals. Then, using the fact that *v*_s_ *+o*_s_ *> v*_p_ *+o*_p_ and that among the surviving offspring there are *n*_s_ scroungers and *n*_p_ producers, solving for 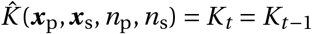 from eq. (C.27) yields,

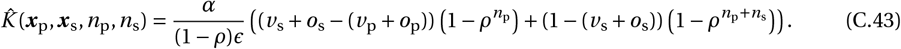

In the large population limit, *n*_p_ *+n*_s_ → ∞, eq. (C.43) converges to

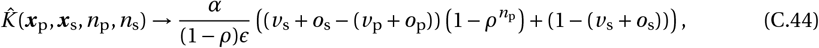

which turns out to be more useful.

Solving eq. (C.40) for 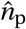 and 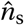 using eqs. (C.41), (C.42) and (C.44) with 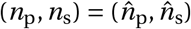, we obtain that

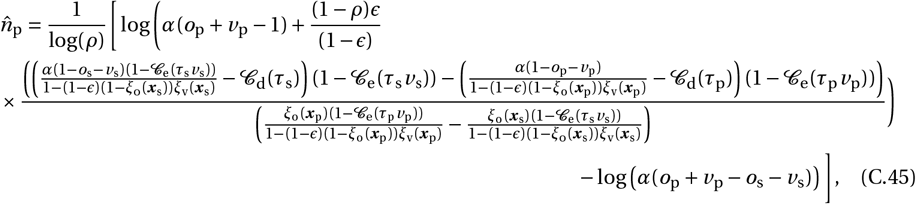

And

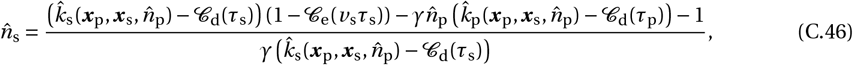

where 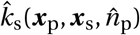 and 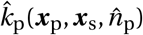 are obtained using eqs. (C.42) and (C.44), for which we removed the dependency on *n*_s_ to highlight that in the large population limit, the equilibrium knowledge within each morph depends on *n*_p_ only.

We computed numerically 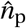 and 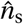 at the early stage of evolutionary branching using eqs. (C.45)–(C.46) for different values of *ρ* and *ϵ*, assuming that the traits of the two morphs ***x***_p_ and ***x***_s_ are given by ***x***_p_ *=* ***x***^*^ *+δ***e**_1_ and ***x***_s_ *=* ***x***^*^ −*δ***e**_1_, where *δ >* 0 is small and **e**_1_ is the normalised eigenvector of **H**(***x***^*^) associated with the leading eigenvalue of **H**(***x***^*^) (with the last component of **e**_1_ positive to ensure that individuals expressing ***x***_p_ perform more teaching than those expressing ***x***_s_). This choice for ***x***_p_ and ***x***_s_ reflects the notion that at an evolutionary branching point, selection is disruptive along the leading eigenvector of the Hessian matrix [26, 27].

Results are shown in Fig. C.1, which reveals that the equilibrium number of producers at the early stage of evolutionary branching is very low for all the parameters explored. This indicates that as soon as polymorphism emerges, selection favours a very small number of producers, which favours their extinction whereby none by chance reproduce. This suggests that the collapse of polymorphism observed in Fig. 4d is due to the stochastic extinction of producers.

**Figure C.1:**
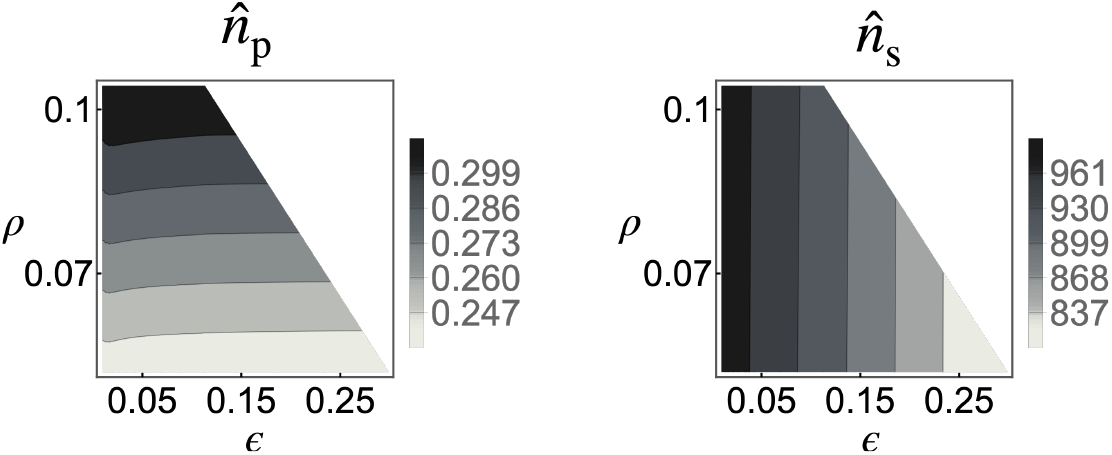
Equilibrium numbers of knowledge producers and scroungers. See eqs. (C.45) and (C.46). Parameters: *δ =* 0.01, *β*_v0_ *=* 7, *β*_*τ*_ *=* 10, *β*_o_ *=* 4, *α =* 2, 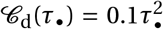, *C*_e_(*v*_*•*_*τ*_*•*_) *=*0.1(*v*_*•*_*τ*_*•*_)^2^, *γ =* 10^−3^.

### D The effect of handling time

Here, we briefly investigate the influence of the time to handle a unit of knowledge *t*_h_ on evolutionary dynamics by looking at how *t*_h_ affects the equilibrium ***x***^*^. To do so, we first consider eq. (A.4) with arbitrary *t*_h_. Using the software Wolfram Mathematica [28], we can obtain an analytical solution for *k*_*•t*_ (1) from eq. (A.4). This solution however is too complicated to be insightful and we therefore refrain from including here, generally writing it as a function *F* of focal traits ***x***_*•*_ and knowledge *k*_*•t*−1_(1) at the previous generation, i.e. as *k*_*•t*_ (1) *= ℱ* (***x***_*•*_, *k*_*•t*−1_(1)). Ideally, we would solve the recurrence defined by this equation for its equilibrium 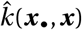, which is such that

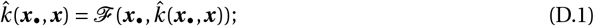

but unfortunately, the function ℱ is too complicated to do so. To circumvent this problem, let us consider the selection gradient on each trait keeping 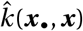 implicit, i.e. we substitute eq. (13) into (14b) keeping 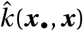 as such, obtaining,

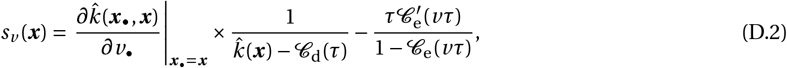

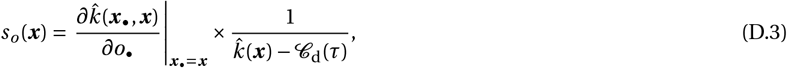

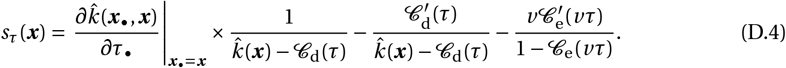

Note that if we substitute eq. (A.19) (which is the solution for 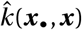 with *t*_h_ *=* 0) into the above, we obtain the selection gradients eqs. (C.1)–(C.3).

To characterise the partial derivative of 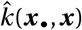 with respect to the focal traits, we differentiate both sides of eq. (D.1) with respect to *z*_*•*_ where *z*_*•*_ ∈ {*v*_*•*_, *o*_*•*_, *τ*_*•*_}, which yields

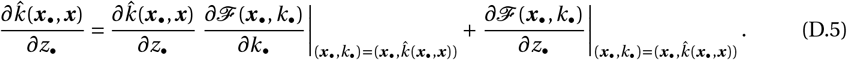

Straightforward re-arrangements lead us to,

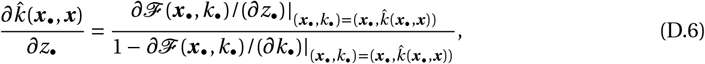

which evaluated at ***x***_*•*_ *=* ***x*** obtains

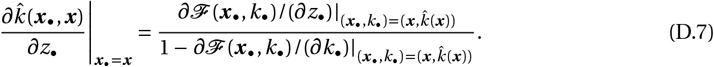

Then, substituting eq. (D.7) into eqs (D.2), (D.3), (D.4), we get

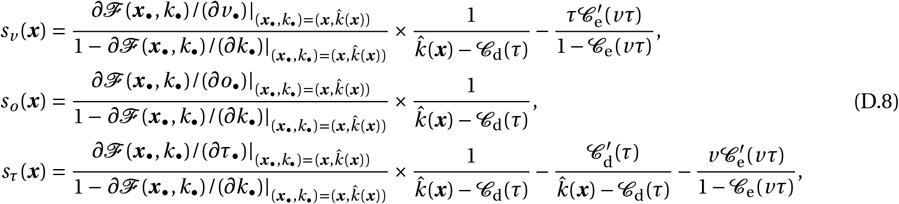

which we use to compute ***x***^*^ numerically following the dynamical procedure described in Appendix C.4.3. Note that 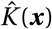, which appears in 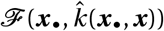, is still given by eq. (A.18), as this approximation re-lies only on the assumption that the population is large and still holds for arbitrary *t*_h_ (as a large population ensures a rapid acquisition of the total available knowledge from the previous generation, irrespective of *t*_h_). Note also that to compute 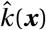, we solved 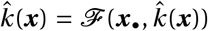 numerically (using the function FindRoot[] in Mathematica). Overall, this numerical procedure is quite heavy and time-consuming. We therefore focused on a limited set of parameters. Our results are shown in Fig. D.1. See main text section 3.2 for interpretation.

**Figure D.1:**
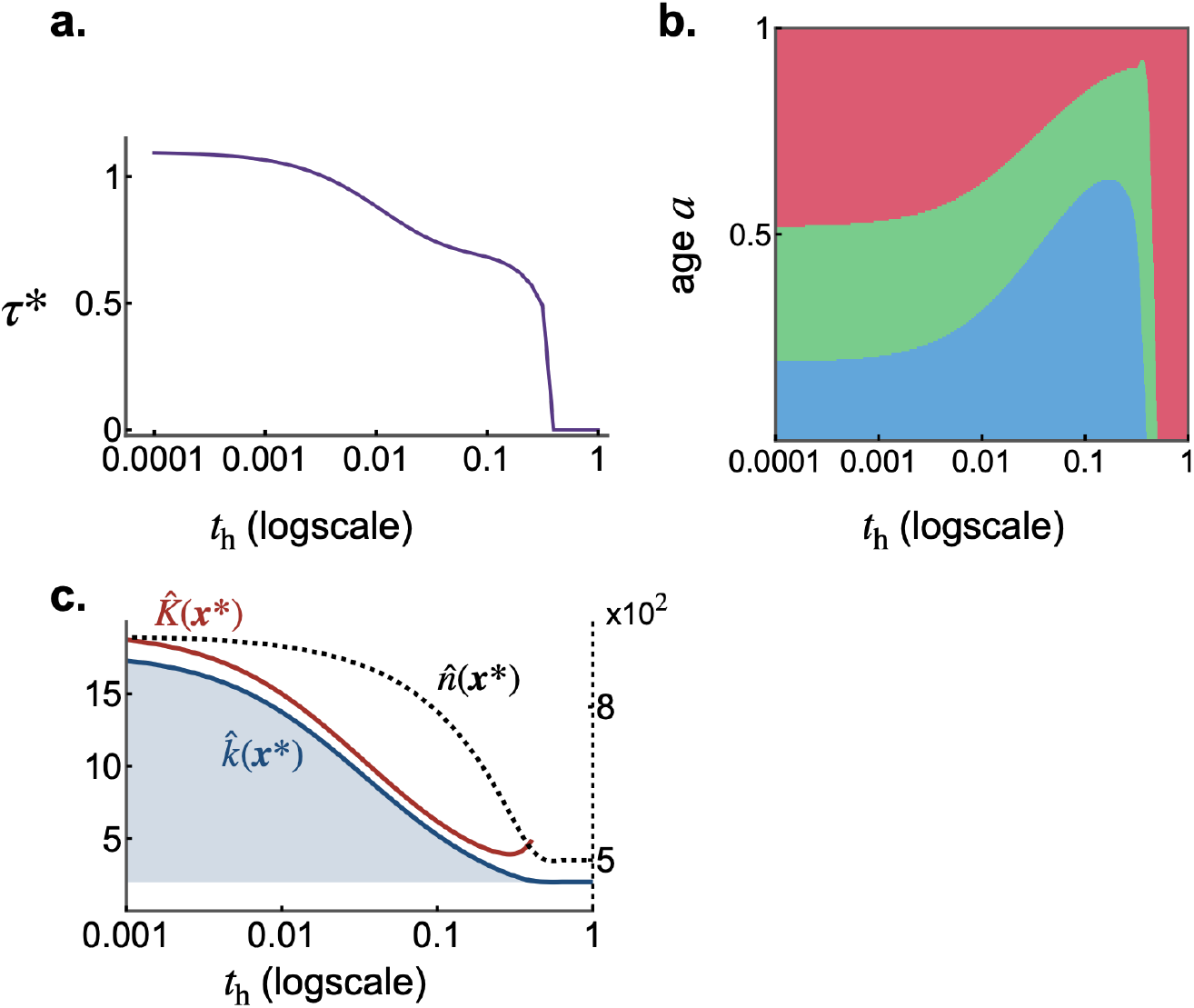
Impact of handling time. *t*_h_ **on the coevolution of learning schedules and teaching. a**. Teaching effort *τ*^*^ at ***x***^*^. **b**. Learning schedule (y-axis) at evolutionary equilibrium ***x***^*^ against *t*_h_ (x-axis, ***x***^*^ computed using eq. D.8). Blue, green, and pink areas represent time spent performing vertical, oblique, and individual learning, respectively. **c**. Individual (obtained by estimating the solution of eq. D.1 with ***x***_*•*_ *=* ***x*** *=* ***x***^*^ using the function FindRoot[] in Mathematica) and total (eq. A.18 with ***x*** *=* ***x***^*^) knowledge, and population size (eq. 12 with ***x*** *=* ***x***^*^) at ***x***^*^. Shaded area corresponds to cumulative knowledge. Left axis gives the scale of knowledge, and right axis the scale of population size. Parameters used in all panels: *β*_v0_ *=* 7, *β*_τ_ *=* 10, *β*_o_ *=* 4, *α =* 2, *ϵ =* 0.1, *ρ =* 0.5, 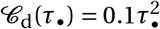, 𝒞_e_(*v*_*•*_*τ*_*•*_) *=* 0.1(*v*_*•*_*τ*_*•*_)^2^, *γ =* 10^−3^.

### E Supplementary discussion

Here we discuss further some of our modelling choices and how departures from these could influence our results. We assumed that the total learning time is fixed, resulting in trade-offs only among the different learning activities. But learning may also take time and energy away from other fitnessrelated activities such as courting, mating, or foraging. We expect that including such costs in our model would not change significantly our results. Individual costs to learning favour lower levels of learning and thus less cumulative knowledge [2, 4, 5, 29–32], but these individual costs can be offset by the indirect fitness benefits of transmitting knowledge to kin when individuals preferentially learn from relatives [2, 7, 33–36]. Such indirect benefits would also be amplified by the evolution of teaching and therefore teaching would also lead to greater levels of cumulative knowledge where learning is individually costly.

Another assumption we made is that teaching boosts vertical transmission by increasing the rate at which offspring encounter novel pieces of knowledge. Alternatively, we could have assumed that individuals need time to handle socially acquired knowledge and that teaching reduces this handling time (i.e. reduce *t*_h_ during vertical learning in eq. A.4). Teaching would also improve the effective learning rate during vertical learning in this case. Evolutionary outcomes should therefore be at least qualitatively similar to the ones we observed here. What may happen though is that the polymorphism between knowledge producers and scroungers shows greater stability where teaching reduces handling time. This is because scroungers would struggle to acquire knowledge obliquely from producers in this case, as this knowledge would be long to handle in the absence of teaching. Presumably, this would result in a more balanced proportion of producers and scroungers and thus lower the risk of scroungers’ extinction.

Finally, our results are based on the assumption that the population size is large. This may favour oblique learning as individuals can learn from a large number of adults, who may thus carry much knowledge. The quantity of knowledge available via oblique learning would tend to be smaller in small populations. This may in turn decrease the time allocated to oblique learning favoured by selection. Nevertheless, the trait means that evolve in individual-based simulations with as few as 1000 individuals match closely the predictions of our quantitative genetics approach (Fig. 3c), which assumes a large population size. This suggests that the effect of population size on evolutionary dynamics is weak, at least provided population size is not extremely small.

## Supplementary figures

**Figure S1:**
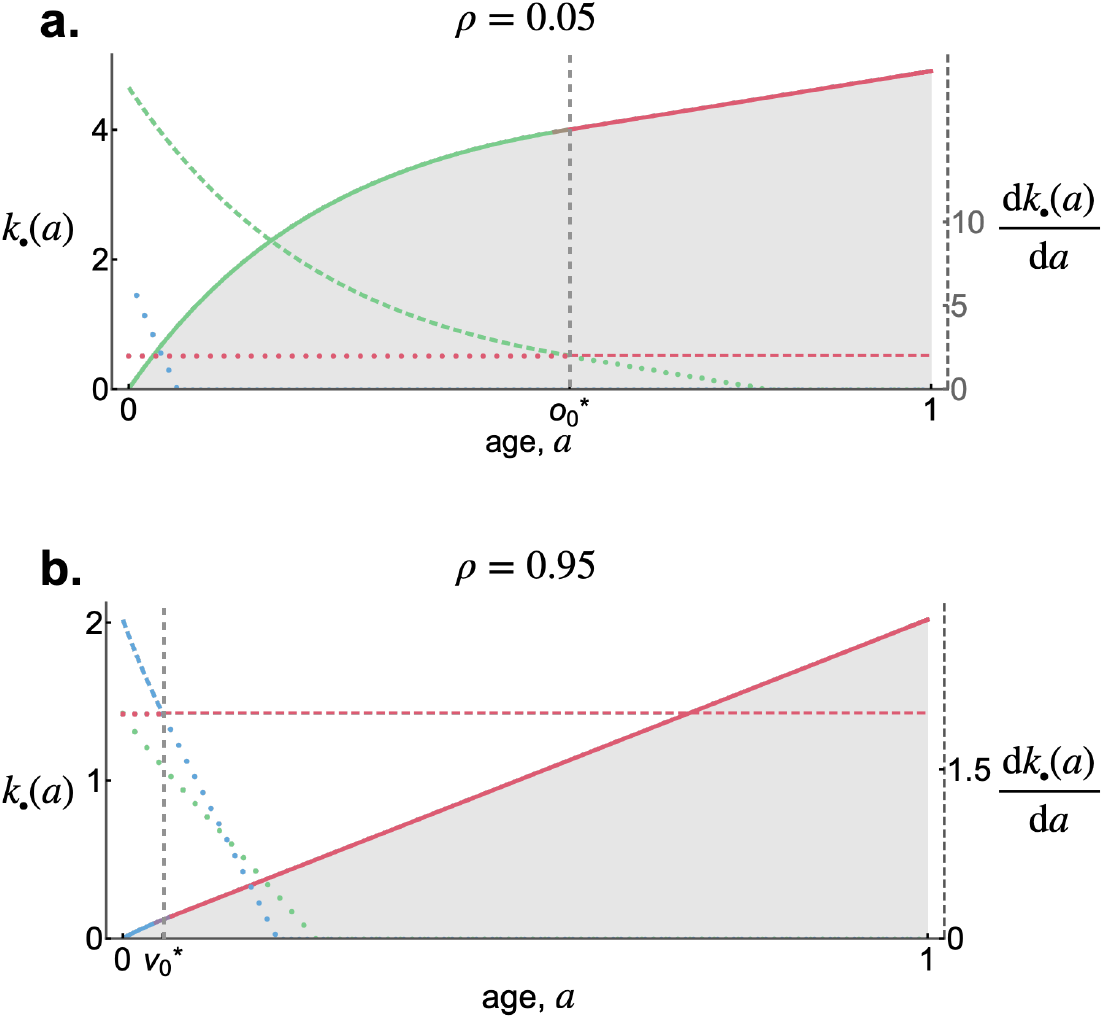
Examples of learning schedules at 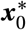 where individuals skip either the vertical or oblique learning phase. Same plot as the main text Fig. 2a. but with **a**. *ρ =* 0.05 and **b**. *ρ =* 0.95. Panels a and b show examples where natural selection promotes skipping the vertical and oblique learning phase, respectively.

**Figure S2:**
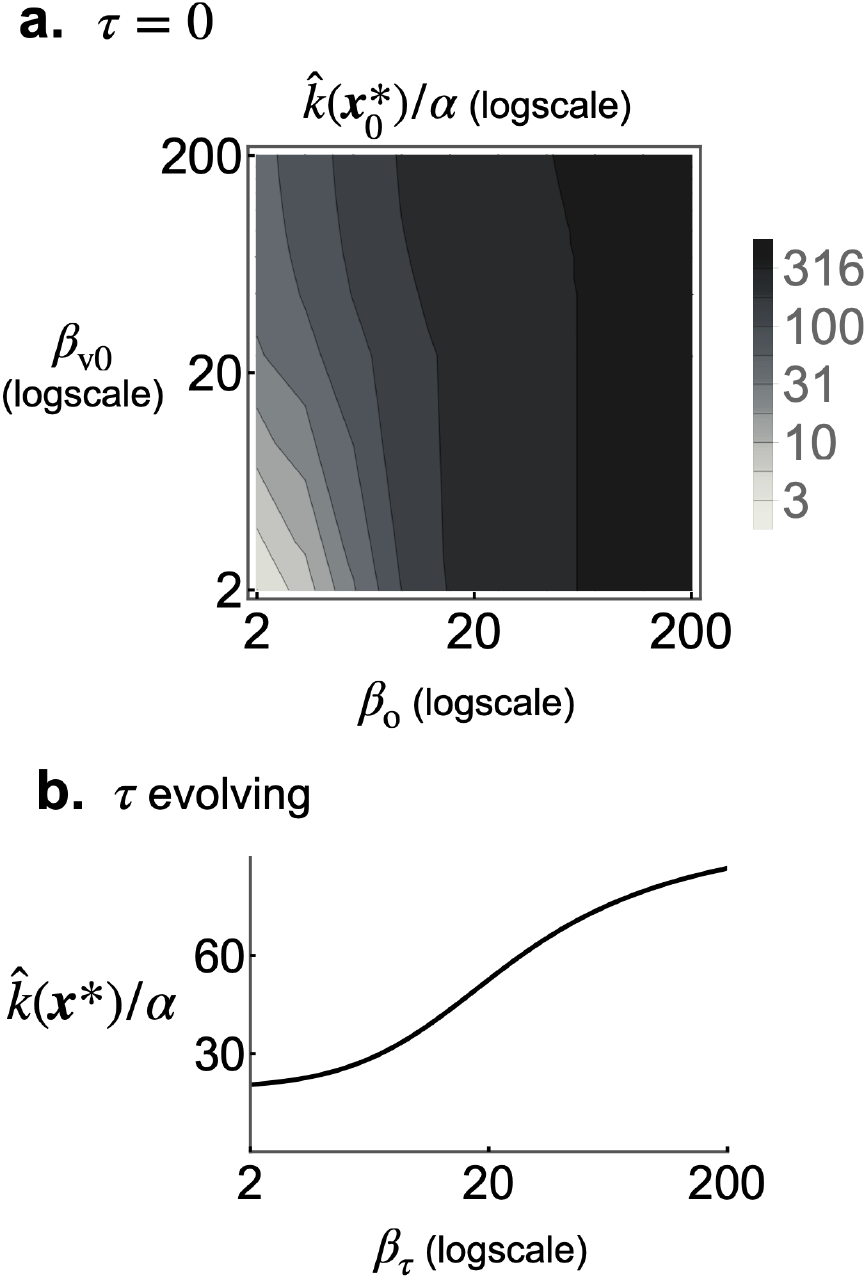
The effect of social learning transmission rates on cumulative knowledge. **a**. Relative individual knowledge 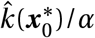 at 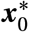 for different values of oblique *β*_o_ and vertical *β*_v0_ transmission rate, shows that cumulative knowledge can become very large where oblique transmission rate *β*_o_ is large. **b**. Relative individual knowledge 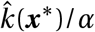 at ***x***^*^ according to teaching efficiency *β*_τ_, showing that greater efficiency boosts cumulative knowledge at evolutionary equilibrium. Parameters: *ϵ =* 0.05, *ρ =* 0.95; other parameters: same as Fig. 3.

**Figure S3:**
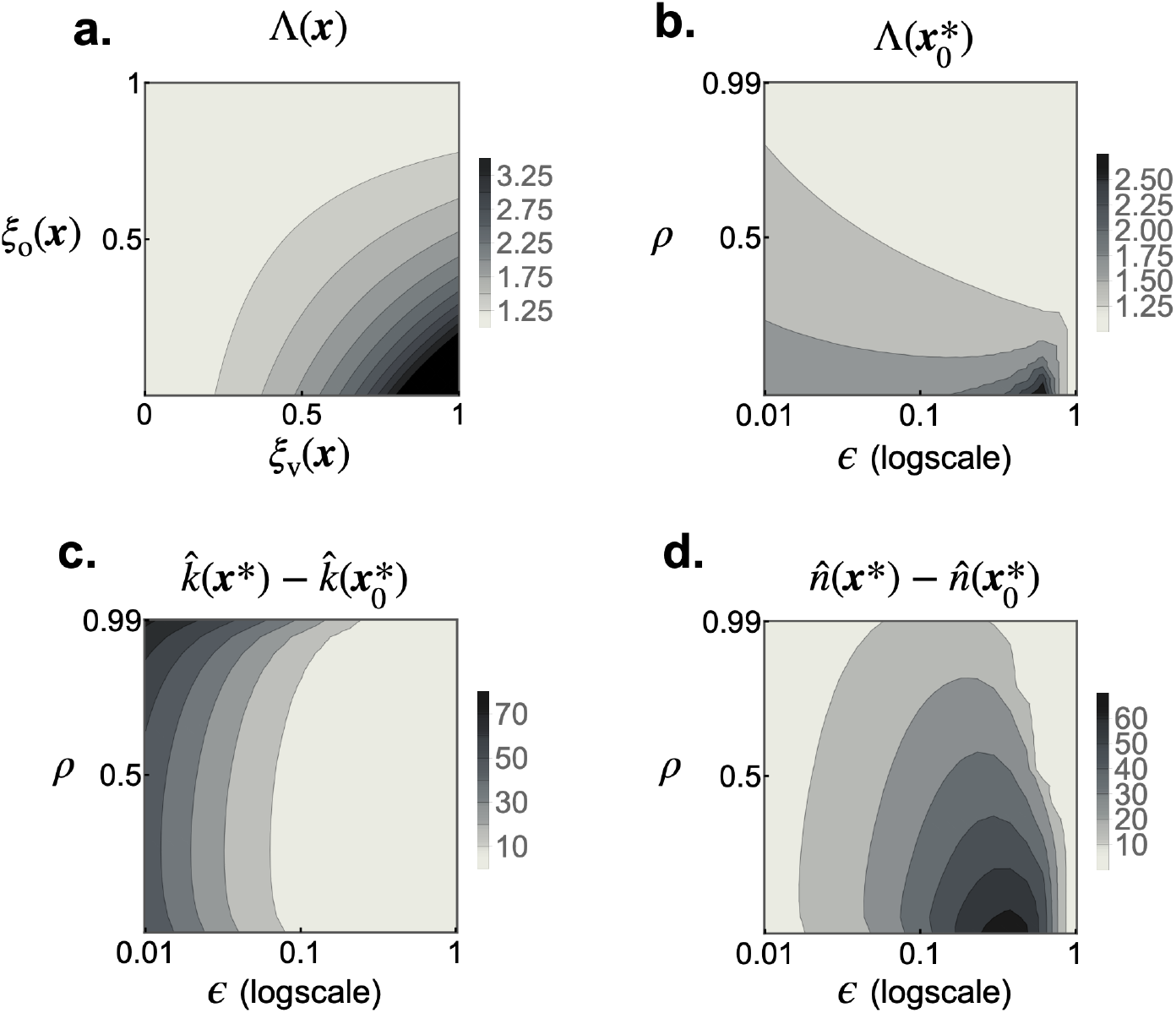
Selection on teaching and its evolutionary impact. **a**. Inter-generational effects of teaching Λ(***x***) as a function of vertical *ξ*_v_(***x***) and oblique *ξ*_o_(***x***) inter-generational transmission rate (with *ϵ =* 0.1). **b**. Inter-generational effects of teaching 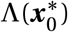 at 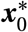 for different values of *ϵ* and *ρ*. **c-d**. The increase in individual knowledge 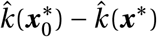 (in panel c) and in the population size 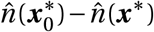 (in panel d) owing to the evolution of teaching for different values of *ρ* and *ϵ*. Parameters: same as in main text Fig. 3.

**Figure S4:**
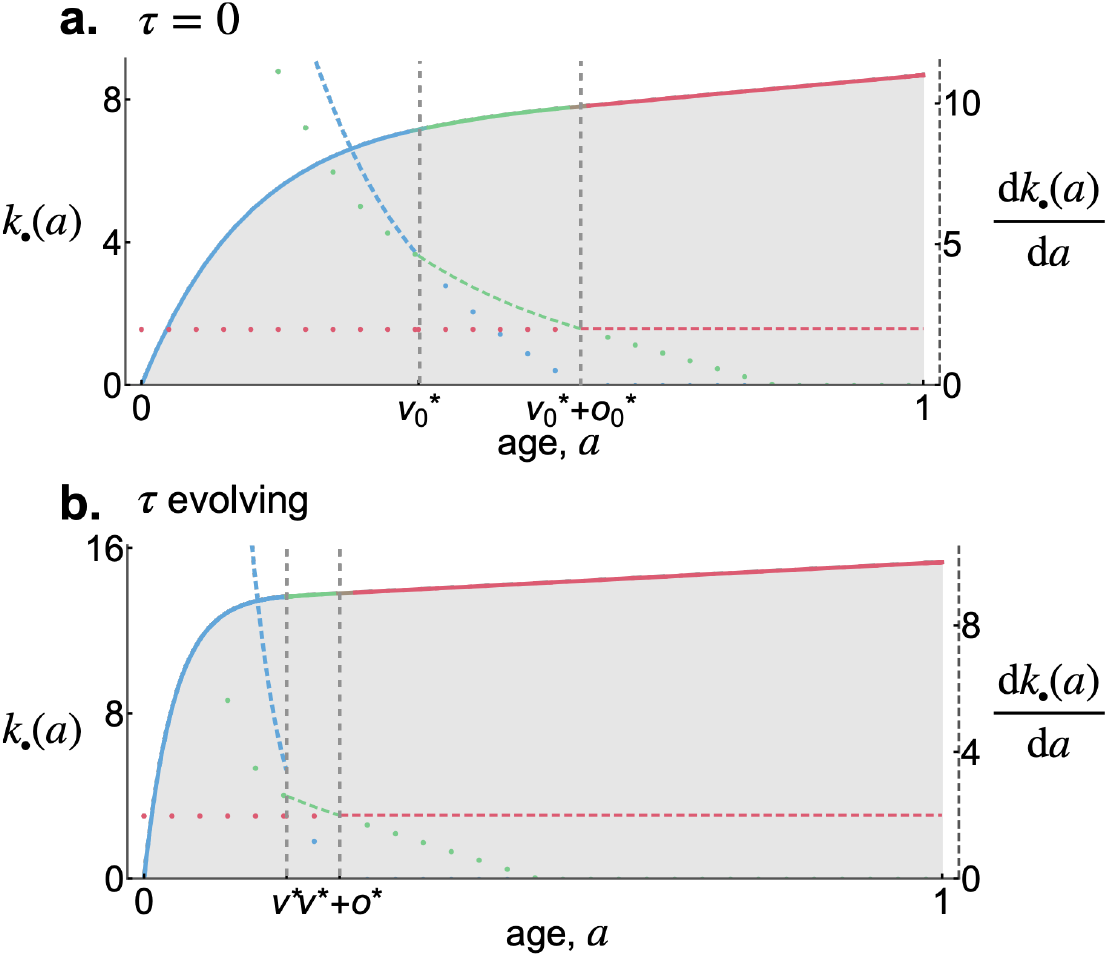
The effect of the evolution of teaching on learning schedule favoured by selection. Panel a shows the learning schedule favoured by selection in the absence of teaching (i.e. at 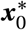 as in main text Fig. 2), while Panel b is the learning schedule at ***x***^*^ where teaching and learning coevolve. Comparing panels a and b shows that the evolution of teaching enables individuals to reach the critical level of knowledge where it becomes more productive to shift to individual learning (where the green dashed and red dashed lines intersect). Parameters: *ϵ =* 0.1, *ρ =* 0.05; other parameters: same as in main text Fig. 3.

**Figure S5:**
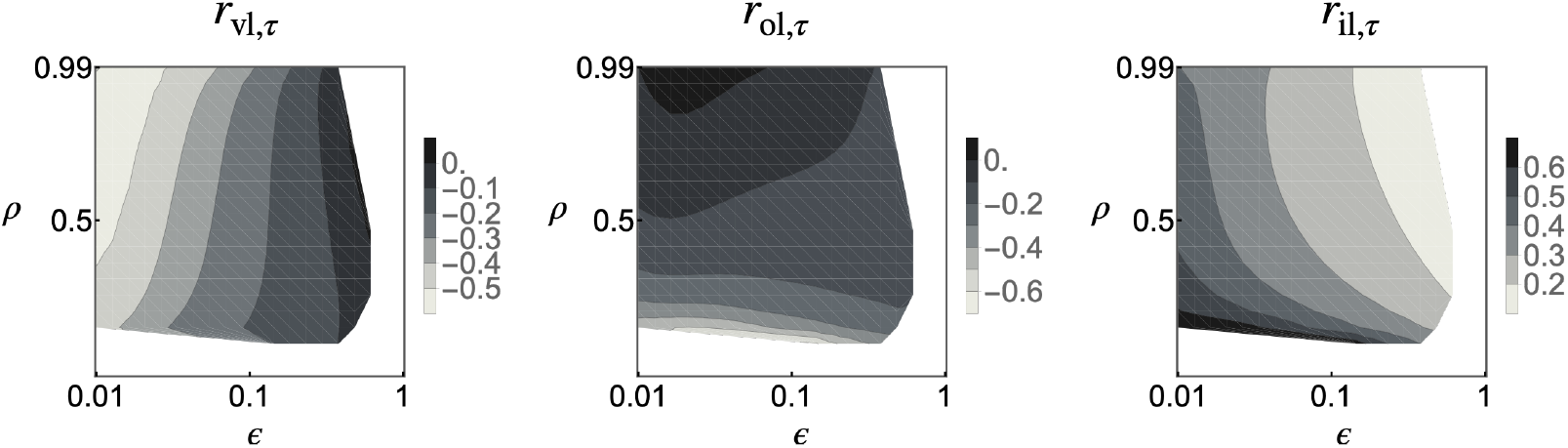
Association between teaching effort and the time spent in each learning phase. Correlation at selection-mutation balance between teaching effort and: (i) vertical learning *r*_vl,*τ*_; (ii) oblique learning *r*_ol,*τ*_; and (iii) individual learning *r*_il,*τ*_ (from eq. B.10 in Appendix). White areas correspond to parameter values where selection is disruptive, or where ***x***^*^ is at the boundary of the phenotypic space. These panels indicate that individuals who teach more tend to spend less time on vertical and oblique learning, and more time on individual learning. Parameters: same as in main text Fig. 3.

**Figure S6:**
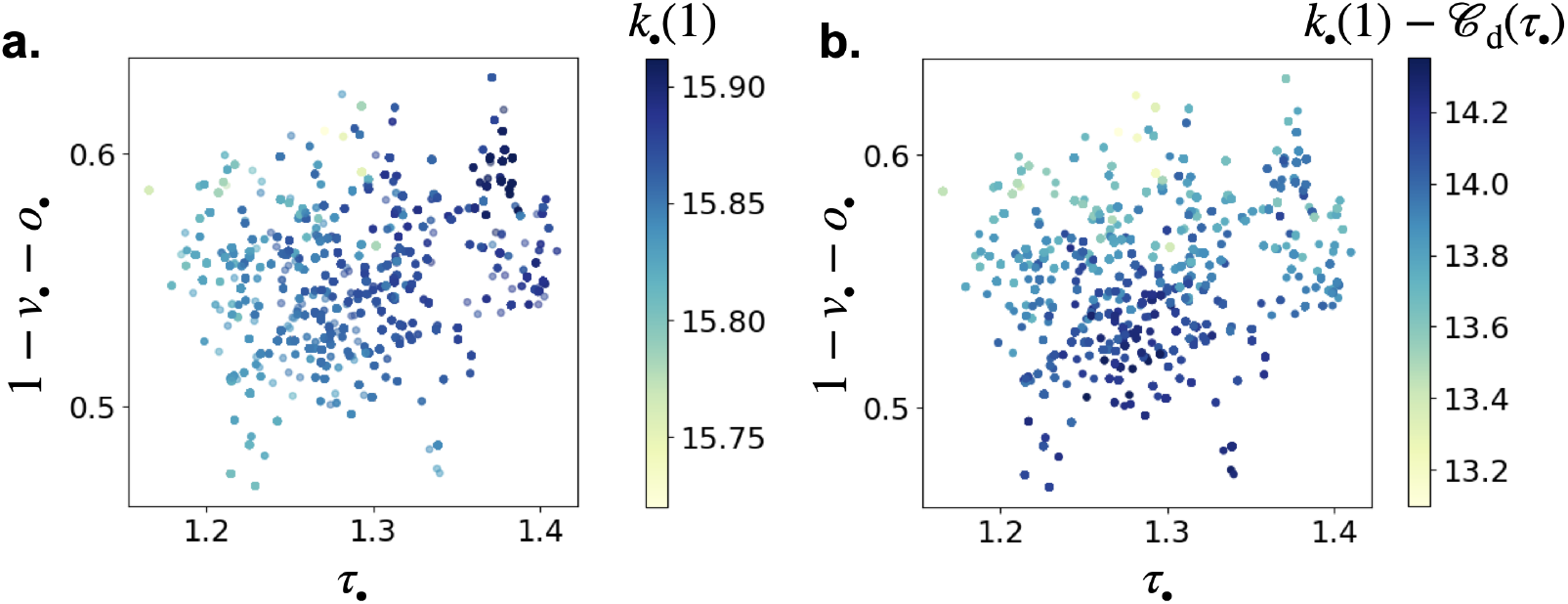
Distribution of knowledge and fecundity according to teaching and individual learning. Snapshot of phenotypic distribution at generation 20, 000 of individual-based simulation shown in Fig. 4b. Colour indicates: **a**. individual knowledge at the age of reproducing *k*_*•*_(1); and **b**. individual fecundity *k*_*•*_(1) − *𝒞*_d_(*τ*_*•*_). Panel a shows that individuals who teach more tend to have greater knowledge (correlation between *τ*_*•*_ and *k*_*•*_(1) is 0.78 here), but Panel b indicates that these individuals do not have higher fertility as teaching incurs fertility costs (correlation between *τ*_*•*_ and fecundity is 0.09 here).

**Figure S7:**
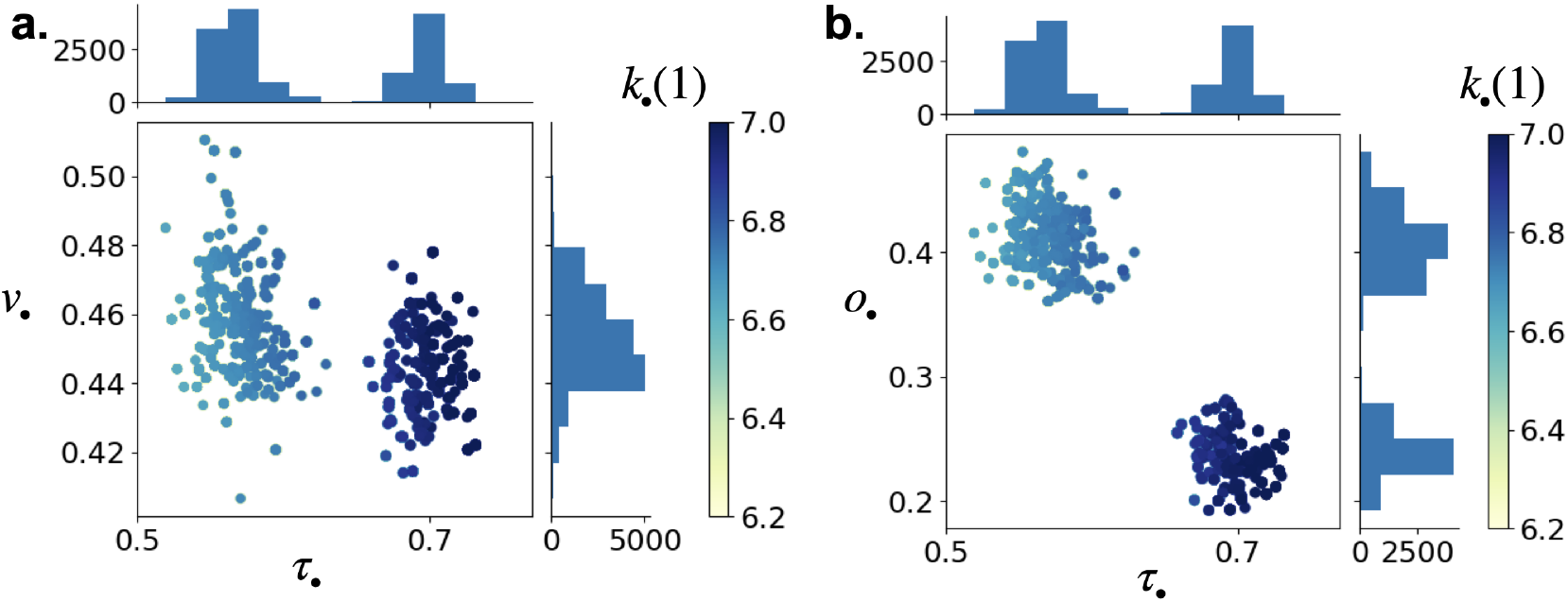
Association between teaching effort and the time spent in vertical and oblique learning phase. Snapshot of phenotypic distribution at generation 7000 of the simulation shown in Fig. 4c. and e. Colour indicates individual knowledge at the age of reproducing *k*_*•*_(1). This shows that knowledge producers (those that teach more) tend to also engage in less vertical learning and less oblique learning than knowledge scroungers.

**Figure S8:**
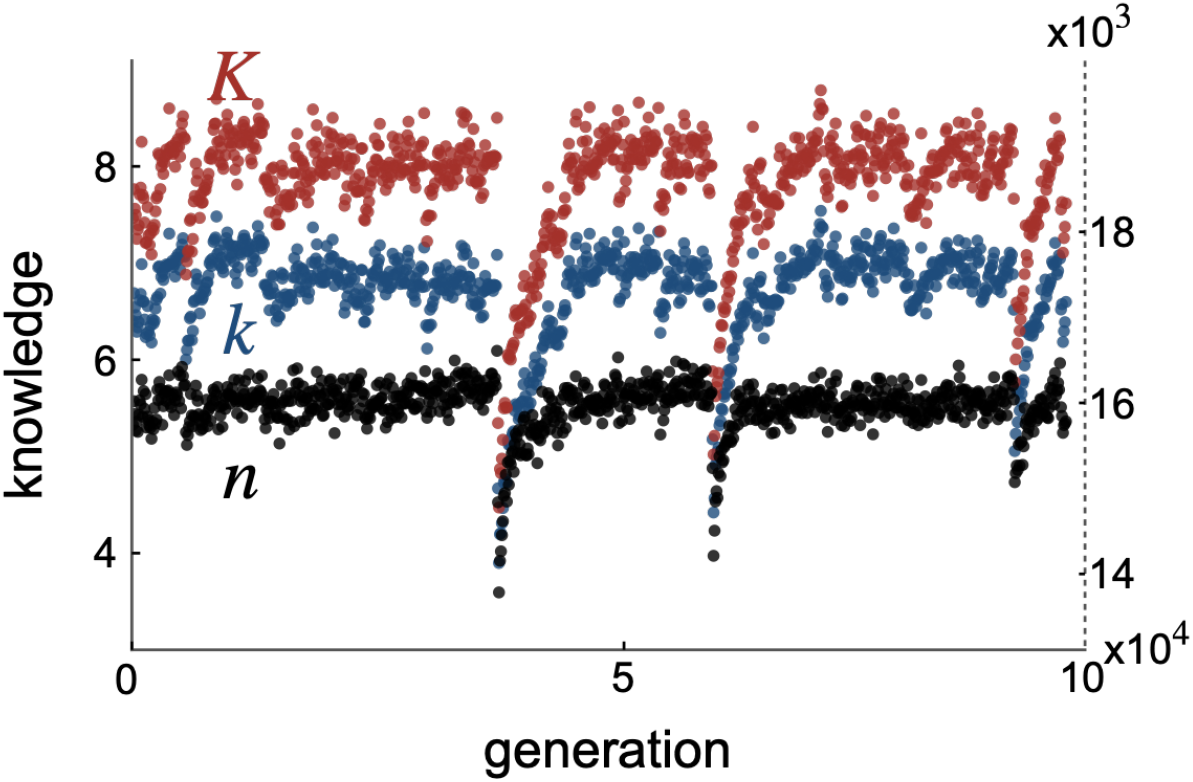
Boom and bust of knowledge and population size. Average individual (blue) and total (red) knowledge in the population whose size is shown in black from same individual based simulation as in Fig. 4d (left axis gives scale of knowledge, and right axis gives scale of population size). This shows that each time the producer morph goes extinct, this results in a sudden drop of knowledge and thus of population size.

## References

[1] L. Dugatkin. Principles of Animal Behavior. W. W. Norton and Company, London, 2004.

[2] E. Thorndike. Animal intelligence: An experimental study of the associative processes in animals. The Psychological Review: Monograph Supplements, 2(4):i, 1898.

[3] T. Zentall. Imitation: definitions, evidence, and mechanisms. Animal Cognition, 9(4):335–353, 2006.

[4] L. Bates and R. Byrne. Imitation: what animal imitation tells us about animal cognition. Wiley interdisciplinary reviews-cognitive science, 1(5):685–695, 2010.

[5] R. Boyd and P. Richerson. An evolutionary model of social learning: The effects of spatial and temporal variation, pages 29–48. Lawrence Erlbaum, 1988.

[6] a. R. Rogers. Does biology constrain culture? American Anthropologist, 90:819–831, 1988.

[7] M. W. Feldman, K. Aoki, and J. Kumm. Individual versus social learning: evolutionary analysis in a fluctuating environment. Anthropological Science, 104:209–231, 1996.

[8] J. Y. Wakano, K. Aoki, and M. W. Feldman. Evolution of social learning: a mathematical analysis. Theoretical Population Biology, 66:249–258, 2004.

[9] P. van den Berg, T. Vu, and L. Molleman. Unpredictable benefits of social information can lead to the evolution of individual differences in social learning. Nature Communications, 15(1):5138, 2024.

[10] L. Lehmann, M. W. Feldman, and R. Kaeuffer. Cumulative cultural dynamics and the coevolution of cultural innovation and transmission: an ESS model for panmictic and structured populations. Journal of Evolutionary Biology, 23:2356–2369, 2010.

[11] L. Rendell, L. Fogarty, and K. N. Laland. Rogers’ paradox recast and resolved: population structure and the evolution of social learning strategies. Evolution, 64(2):534–548, 2010.

[12] H. Ohtsuki, J. Y. Wakano, and Y. Kobayashi. Inclusive fitness analysis of cumulative cultural evolution in an island-structured population. Theoretical Population Biology, 2017.

[13] M. Ammar, L. Fogarty, and A. Kandler. Social learning and memory. Proceedings of the National Academy of Sciences, 120(33):e2310033120, 2023.

[14] K. Aoki, J. Y. Wakano, and L. Lehmann. Evolutionarily stable learning schedules and cumulative culture in discrete generation models. Theoretical Population Biology, 81(4):300–309, 2012.

[15] L. Lehmann, J. Y. Wakano, and K. Aoki. On optimal learning schedules and the marginal value of cumulative cultural evolution. Evolution, 67:1435–1445, 2013.

[16] J. Y. Wakano and C. Miura. Trade-off between learning and exploitation: The Pareto-optimal versus evolutionarily stable learning schedule in cumulative cultural evolution. Theoretical Population Biology, 91:37–43, 2014.

[17] J.-B. André and N. Baumard. Cultural evolution by capital accumulation. Evolutionary Human Sciences, 2:1–13, 2020.

[18] C. P. van Schaik. The Primate Origin of Human Behavior. Wiley-Blackwell, New Jersey, 2016.

[19] D. Acemoglu. Introduction to Modern Economic Growth. Princeton University Press, Princeton, NJ, 2009.

[20] G. R. Hunt and R. D. Gray. Diversification and cumulative evolution in new caledonian crow tool manufacture. Proceedings of the Royal Society of London. Series B: Biological Sciences, 270 (1517):867–874, 2003.

[21] T. Sasaki and D. Biro. Cumulative culture can emerge from collective intelligence in animal groups. Nature Communications, 8(1):15049, 2017.

[22] B. R. Jesmer, J. A. Merkle, J. R. Goheen, E. O. Aikens, J. L. Beck, A. B. Courtemanch, M. A. Hurley, D. E. McWhirter, H. M. Miyasaki, K. L. Monteith, and M. J. Kauffman. Is ungulate migration culturally transmitted? evidence of social learning from translocated animals. Science, 361(6406): 1023–1025, 2018.

[23] M. Enquist, K. Eriksson, and S. Ghirlanda. Critical social learning: a solution to Rogers’s paradox of nonadaptive culture. American Anthropologist, 109:727–734, 2007.

[24] K. Aoki. Evolution of the social-learner-explorer strategy in an environmentally heterogeneous two-island model. Evolution, 64(9):2575–2586, 2010.

[25] Y. Kobayashi, H. Ohtsuki, and J. Y. Wakano. Population size vs. social connectedness — a gene-culture coevolutionary approach to cumulative cultural evolution. Theoretical Population Biology, 111:87–95, 2016.

[26] Y. Kobayashi, J. Y. Wakano, and H. Ohtsuki. A paradox of cumulative culture. Journal of Theoretical Biology, 379:79–88, 2015.

[27] C. Mullon and L. Lehmann. Invasion fitness for gene-culture co-evolution in family-structured populations and an application to cumulative culture under vertical transmission. Theoretical Population Biology, 116:33–46, 2017.

[28] Y. Kobayashi, J. Y. Wakano, and H. Ohtsuki. Evolution of cumulative culture for niche construction. Journal of Theoretical Biology, 472:67–76, 2019.

[29] C. Mullon, J. Y. Wakano, and H. Ohtsuki. Coevolutionary dynamics of genetic traits and their long-term extended effects under non-random interactions. Journal of Theoretical Biology, 525: 110750, 2021.

[30] L. Fogarty, P. Strimling, and K. N. Laland. The evolution of teaching. Evolution, 65(10):2760–2770, 2011.

[31] L. Castro and M. A. Toro. Cumulative cultural evolution: The role of teaching. Journal of Theoretical Biology, 347:74–83, 2014.

[32] N. R. Franks and T. Richardson. Teaching in tandem-running ants. Nature, 439(7073):153–153, 2006.

[33] A. Thornton and K. McAuliffe. Teaching in wild meerkats. Science, 313(5784):227–229, 2006.

[34] A. H. Boyette and B. S. Hewlett. Teaching in hunter-gatherers. Review of Philosophy and Psychology, 9(4):771–797, 2018.

[35] C. Boesch. Teaching among wild chimpanzees. Animal Behaviour, 41(3):530–532, 1991.

[36] D. Maestripieri. First steps in the macaque world: do rhesus mothers encourage their infants’ independent locomotion? Animal Behaviour, 49(6):1541–1549, 1995.

[37] D. Maestripieri. Maternal encouragement of infant locomotion in pigtail macaques,macaca nemestrina. Animal Behaviour, 51(3):603–610, 1996.

[38] N. J. Raihani and A. R. Ridley. Experimental evidence for teaching in wild pied babblers. Animal Behaviour, 75(1):3–11, 2008.

[39] L. G. Rapaport. Progressive parenting behavior in wild golden lion tamarins. Behavioral Ecology, 22(4):745–754, 2011.

[40] S. Kleindorfer, H. Hoi, C. Evans, K. Mahr, J. Robertson, M. Hauber, and D. Colombelli-Négrel. The cost of teaching embryos in superb fairy-wrens. Behavioral Ecology, 25:1131–1135, 2014.

[41] S. Musgrave, D. Morgan, E. Lonsdorf, R. Mundry, and C. Sanz. Tool transfers are a form of teaching among chimpanzees. Scientific Reports, 6(1):34783, 2016.

[42] C. A. Troisi, W. J. E. Hoppitt, C. R. Ruiz-Miranda, and K. N. Laland. Food-offering calls in wild golden lion tamarins (leontopithecus rosalia): Evidence for teaching behavior? International Journal of Primatology, 39(6):1105–1123, 2018.

[43] T. M. Caro and M. D. Hauser. Is there teaching in nonhuman animals? The quarterly review of biology, 67(2):151–174, 1992.

[44] Hoppitt, W. J. E, G. R. Brown, R. Kendal, L. Rendell, A. Thornton, M. M. Webster, and K. N. Laland. Lessons from animal teaching. Trends in Ecology and Evolution, 23:486–493, 2008.

[45] A. Thornton and N. J. Raihani. The evolution of teaching. Animal Behaviour, 75:1823–1836, 2008.

[46] M. A. Kline. How to learn about teaching: An evolutionary framework for the study of teaching behavior in humans and other animals. Behavioral and Brain sciences, 38:e31, 2015.

[47] D. Frye and M. Ziv. Teaching and learning as intentional activities. In The development of social cognition and communication, pages 231–258. Psychology Press, 2013.

[48] S. Strauss, M. Ziv, and D. Frye. Cognitive universals and cultural variation in teaching. Behavioral and Brain Sciences, 38:e67, 2015.

[49] L. Castro, M. A. Toro, et al. Cultural transmission and the capacity to approve or disapprove of offspring’s behaviour. Journal of Memetics-Evolutionary Models of Information Transmission, 6 (2):1–17, 2002.

[50] K. Aoki, J. Y. Wakano, and M. W. Feldman. Gene-culture models for the evolution of altruistic teaching. In M. Tibayrenc and F. Ayala, editors, On human nature: biology, psychology, ethics, politics, and religion, pages 279–296. Academic Press. Amsterdam (The Netherlands) and Boston (Massachusetts): Elsevier, 2017.

[51] M. D. Gurven, R. J. Davison, and T. S. Kraft. The optimal timing of teaching and learning across the life course. Philosophical Transactions of the Royal Society B: Biological Sciences, 375(1803): 20190500, 2020.

[52] R. Ventura and E. Akcay. A cognitive-evolutionary model for the evolution of teaching. Journal of Theoretical Biology, 533:110933, 2022.

[53] C. R. Turner, S. F. Mann, M. Spike, R. D. Magrath, and K. Sterelny. Joint evolution of traits for social learning. Behavioral Ecology and Sociobiology, 77(4):47, 2023.

[54] H. Kaplan, J. Lancaster, and A. Robson. Embodied capital and the evolutionary economics of the human life span. In J. Carey and S. Tuljapurkar, editors, Life span: evolutionary, ecological and demographic perspectives, pages 152–182. Univ Calif Berkeley, Ctr Econ & Demog Aging; Natl Inst Aging, 2003. Conference on Life Span - Evolutionary, Ecological, and Demographic Perspectives, Santorini, Greece, May 14-18, 2001.

[55] R. J. Beverton and S. J. Holt. On the dynamics of exploited fish populations. Springer Dordrecht, 1957.

[56] C. Mullon and L. Lehmann. An evolutionary quantitative genetics model for phenotypic (co)variances under limited dispersal, with an application to socially synergistic traits. Evolution, 73(9):1695–1728, 2019.

[57] A. Sih, A. M. Bell, J. C. Johnson, and R. E. Ziemba. Behavioral syndromes: An integrative overview. The Quarterly Review of Biology, 79(3):241–277, 2004. PMID: 15529965.

[58] A. Sih, A. Bell, and J. Johnson. Behavioral syndromes: an ecological and evolutionary overview. Trends in Ecology & Evolution, 19(7):372–378, 2004.

[59] T. Caraco and L.-A. Giraldea. Social foraging: Producing and scrounging in a stochastic environment. Journal of Theoretical Biology, 153(4):559–583, 1991.

[60] W. L. Vickery, L.-A. Giraldeau, J. J. Templeton, D. L. Kramer, and C. A. Chapman. Producers, scroungers, and group foraging. The American Naturalist, 137(6):847–863, 1991.

[61] G. Beauchamp and L.-A. Giraldeau. Group foraging revisited: information sharing or producer-scrounger game? The American Naturalist, 148(4):738–743, 1996.

[62] I. M. Hamilton. Kleptoparasitism and the distribution of unequal competitors. Behavioral Ecology, 13(2):260–267, 2002.

[63] G. Beauchamp. A spatial model of producing and scrounging. Animal Behaviour, 76(6):1935–1942, 2008.

[64] C. Tennie, J. Call, and M. Tomasello. Ratcheting up the ratchet: on the evolution of cumulative culture. Philosophical transactions of the Royal Society of London. Series B, Biological sciences, 364(1528):2405–2415, 2009.

[65] H. M. Lewis and K. N. Laland. Transmission fidelity is the key to the build-up of cumulative culture. Philosophical Transactions of the Royal Society B: Biological Sciences, 367(1599):2171–2180, 2012.

[66] R. McElreath and P. Strimling. When natural selection favors imitation of parents. Current Anthropology, 49:307–316, 2008.

[67] R. Byrne and A. Whiten. Tactical deception of familiar individuals in baboons (papio ursinus). Animal Behaviour, 33(2):669–673, 1985.

[68] C. Marchetti and P. J. Drent. Individual differences in the use of social information in foraging by captive great tits. Animal Behaviour, 60(1):131–140, 2000.

[69] J. Bouchard, W. Goodyer, and L. Lefebvre. Social learning and innovation are positively correlated in pigeons (columba livia). Animal Cognition, 10(2):259–266, 2007.

[70] R. H. J. M. Kurvers, K. Van Oers, B. A. Nolet, R. M. Jonker, S. E. Van Wieren, H. H. T. Prins, and R. C. Ydenberg. Personality predicts the use of social information. Ecology Letters, 13(7):829–837, 2010.

[71] E. Katsnelson, U. Motro, M. W. Feldman, and A. Lotem. Individual-learning ability predicts social-foraging strategy in house sparrows. Proceedings of the Royal Society B: Biological Sciences, 278(1705):582–589, 2011.

[72] P. Rosa, V. Nguyen, and F. Dubois. Individual differences in sampling behaviour predict social information use in zebra finches. Behavioral Ecology and Sociobiology, 66(9):1259–1265, 2012.

[73] I. Gavriilidi, S. Baeckens, G. De Meester, L. Van Linden, and R. Van Damme. The gullible genius: fast learners fall for fake news. Behavioral Ecology and Sociobiology, 76(1):5, 2021.

[74] S. Nomakuchi, P. J. Park, and M. A. Bell. Correlation between exploration activity and use of social information in three-spined sticklebacks. Behavioral Ecology, 20(2):340–345, 2009.

[75] J. Foucaud, A.-S. Philippe, C. Moreno, and F. Mery. A genetic polymorphism affecting reliance on personal versus public information in a spatial learning task in Drosophila melanogaster. Proceedings of the Royal Society B: Biological Sciences, 280(1760):20130588, 2013.

[76] J. M. Burkart, A. Strasser, and M. Foglia. Trade-offs between social learning and individual innovativeness in common marmosets, callithrix jacchus. Animal Behaviour, 77(5):1291–1301, 2009.

[77] A. Mesoudi. An experimental comparison of human social learning strategies: payoff-biased social learning is adaptive but underused. Evolution and Human Behavior, 32(5):334–342, 2011.

[78] R. Croston, C. Branch, D. Kozlovsky, R. Dukas, and V. Pravosudov. Heritability and the evolution of cognitive traits. Behavioral Ecology, 26(6):1447–1459, 2015.

[79] A. Thornton and J. Samson. Innovative problem solving in wild meerkats. Animal Behaviour, 83 (6):1459–1468, 2012.

[80] M. Kline, R. Boyd, and J. Henrich. Teaching and the life history of cultural transmission in fijian villages. Human Nature, in press, 2013.

[81] A. Thornton and K. McAuliffe. Teaching in wild meerkats. Science, 313(5784):227–229, 2006.

[82] C. Loverdo and H. Viciana. Cultural transmission and biological markets. Biology & Philosophy, 33(5):40, 2018.

[83] J. Henrich and F. J. Gil-White. The evolution of prestige: freely conferred deference as a mechanism for enhancing the benefits of cultural transmission. Evolution and Human Behavior, 22 (3):165–196, 2001.

[84] B. Sinervo and E. Svensson. Correlational selection and the evolution of genomic architecture. Heredity (Edinb)., 89(5):329–338, 2002.

[85] S. M. Reader and K. N. Laland. Primate innovation: sex, age and social rank differences. Internation Journal of Primatology, 22:787–805, 2001.

[86] J. C. Biesmeijer and T. D. Seeley. The use of waggle dance information by honey bees throughout their foraging careers. Behavioral Ecology and Sociobiology, 59:133–142, 2005.

[87] D. W. A. Noble, R. W. Byrne, and M. J. Whiting. Age-dependent social learning in a lizard. Biology Letters, 10(7), 2014.

[88] K. Carr, R. L. Kendal, and E. G. Flynn. Imitate or innovate? children’s innovation is influenced by the efficacy of observed behaviour. Cognition, 142:322–332, 2015.

[89] B. S. Hewlett, H. N. Fouts, A. H. Boyette, and B. L. Hewlett. Social learning among Congo Basin hunter-gatherers. Philosophical transactions of the Royal Society of London. Series B, Biological sciences, 366(1567):1168–1178, 2011.

[90] K. Demps, F. Zorondo-Rodríguez, C. García, and V. Reyes-García. Social learning across the life cycle: cultural knowledge acquisition for honey collection among the Jenu Kuruba India. Evolution and Human Behavior, 33:460–470, 2012.

[91] R. Lande. Quantitative Genetic Analysis of Multivariate Evolution, Applied to Brain: Body Size Allometry. Evolution, 33(1):402, 1979.

[92] J. Y. Wakano and Y. Iwasa. Evolutionary branching in a finite population: deterministic branching vs. stochastic branching. Genetics, 193(1):229–241, 2013.

[93] F. Débarre, S. L. Nuismer, and M. Doebeli. Multidimensional (co)evolutionary stability. Am. Nat., 184(2):158–171, 2014.

[94] J. Y. Wakano and L. Lehmann. Evolutionary branching in deme-structured populations. Journal of Theoretical Biology, 351:83–95, 2014.

[95] F. Débarre and S. P. Otto. Evolutionary dynamics of a quantitative trait in a finite asexual population. Theoretical Population Biology, 108:75–88, 2016.

[96] S. A. H. Geritz, J. A. J. Metz, and C. Rueffler. Mutual invadability near evolutionarily singular strategies for multivariate traits, with special reference to the strongly convergence stable case. Journal of Mathematical Biology, 72(4):1081–1099, 2016.

[97] C. Mullon, L. Keller, and L. Lehmann. Evolutionary stability of jointly evolving traits in subdivided populations. American Naturalist, 188:175–195, 2016.

## References

[1] D. W. Stephens and J. R. Krebs. Foraging Theory, volume 1. Princeton University Press, 1986.

[2] L. Lehmann, M. W. Feldman, and R. Kaeuffer. Cumulative cultural dynamics and the coevolution of cultural innovation and transmission: an ESS model for panmictic and structured populations. Journal of Evolutionary Biology, 23:2356–2369, 2010.

[3] K. Aoki, J. Y. Wakano, and L. Lehmann. Evolutionarily stable learning schedules and cumulative culture in discrete generation models. Theoretical Population Biology, 81(4):300–309, 2012.

[4] L. Lehmann, J. Y. Wakano, and K. Aoki. On optimal learning schedules and the marginal value of cumulative cultural evolution. Evolution, 67:1435–1445, 2013.

[5] J. Y. Wakano and C. Miura. Trade-off between learning and exploitation: The Pareto-optimal versus evolutionarily stable learning schedule in cumulative cultural evolution. Theoretical Population Biology, 91:37–43, 2014.

[6] Y. Kobayashi, H. Ohtsuki, and J. Y. Wakano. Population size vs. social connectedness — a gene-culture coevolutionary approach to cumulative cultural evolution. Theoretical Population Biology, 111:87–95, 2016.

[7] C. Mullon and L. Lehmann. Invasion fitness for gene-culture co-evolution in family-structured populations and an application to cumulative culture under vertical transmission. Theoretical Population Biology, 116:33–46, 2017.

[8] J.-B. André and N. Baumard. Cultural evolution by capital accumulation. Evolutionary Human Sciences, 2:1–13, 2020.

[9] J. Y. Wakano and Y. Iwasa. Evolutionary branching in a finite population: deterministic branching vs. stochastic branching. Genetics, 193(1):229–241, 2013.

[10] F. Débarre, S. L. Nuismer, and M. Doebeli. Multidimensional (co)evolutionary stability. Am. Nat., 184(2):158–171, 2014.

[11] J. Y. Wakano and L. Lehmann. Evolutionary branching in deme-structured populations. Journal of Theoretical Biology, 351:83–95, 2014.

[12] F. Débarre and S. P. Otto. Evolutionary dynamics of a quantitative trait in a finite asexual population. Theoretical Population Biology, 108:75–88, 2016.

[13] C. Mullon and L. Lehmann. An evolutionary quantitative genetics model for phenotypic (co)variances under limited dispersal, with an application to socially synergistic traits. Evolution, 73(9):1695–1728, 2019.

[14] Y. Iwasa, A. Pomiankowski, and S. Nee. The evolution of costly mate preferences II. The ‘handicap’ principle. Evolution, 45:1431–1442, 1991.

[15] M. L. Taper and T. J. Case. Models of character displacement adn the theoretical robustness of taxon cycles. Evolution, 46:317–333, 1992.

[16] P. A. Abrams, Y. Harada, and H. Matsuda. On the Relationship Between Quantitative Genetic and Ess Models. Evolution (N. Y)., 47(3):982–985, 1993.

[17] P. A. Abrams, H. Matsuda, and Y. Harada. Evolutionarily unstable fitness maxima and stable fitness minima of continuous traits. Evol. Ecol., 7(5):465–487, 1993.

[18] S. A. H. Geritz, É. Kisdi, G. Meszéna, and J. A. J. Metz. Evolutionarily singular strategies and the adaptive growth and branching of the evolutionary tree. Evol. Ecol., 12(1):35–57, 1998.

[19] F. Rousset. Genetic Structure and Selection in Subdivided Populations. Princeton University Press, Princeton, NJ, 2004.

[20] F. Dercole and S. Rinaldi. Analysis of Evolutionary Processes: The Adaptive Dynamics Approach and Its Applications. Princeton University Press, Princeton, NJ, 2008.

[21] O. Leimar. Multidimensional convergence stability. Evolutionary Ecology Research, 11(2):191–208, 2009.

[22] S. A. H. Geritz, J. A. J. Metz, and C. Rueffler. Mutual invadability near evolutionarily singular strategies for multivariate traits, with special reference to the strongly convergence stable case. Journal of Mathematical Biology, 72(4):1081–1099, 2016.

[23] R. Lande and S. J. Arnold. The Measurement of Selection on Correlated Characters. Evolution, 37(6):1210–1226, 1983.

[24] A. Sasaki and U. Dieckmann. Oligomorphic dynamics for analyzing the quantitative genetics of adaptive speciation. J. Math. Biol., 63(4):601–635, 2011.

[25] S. Lion, A. Sasaki, and M. Boots. Extending eco-evolutionary theory with oligomorphic dynamics. Ecology Letters, 26(S1):S22–S46, 2023.

[26] C. Mullon, L. Keller, and L. Lehmann. Evolutionary stability of jointly evolving traits in subdivided populations. American Naturalist, 188:175–195, 2016.

[27] S. A. H. Geritz, J. a. J. Metz, and C. Rueffler. Mutual invadability near evolutionarily singular strategies for multivariate traits, with special reference to the strongly convergence stable case. J. Math. Biol., 72(4):1081–1099, 2016.

[28] Wolfram Research, Inc. Mathematica. Wolfram Research, Inc., Champaign, Illinois, 2016.

[29] R. Boyd and P. Richerson. An evolutionary model of social learning: The effects of spatial and temporal variation, pages 29–48. Lawrence Erlbaum, 1988.

[30] a. R. Rogers. Does biology constrain culture? American Anthropologist, 90:819–831, 1988.

[31] M. W. Feldman, K. Aoki, and J. Kumm. Individual versus social learning: evolutionary analysis in a fluctuating environment. Anthropological Science, 104:209–231, 1996.

[32] J. Y. Wakano, K. Aoki, and M. W. Feldman. Evolution of social learning: a mathematical analysis. Theoretical Population Biology, 66:249–258, 2004.

[33] Y. Kobayashi, J. Y. Wakano, and H. Ohtsuki. A paradox of cumulative culture. Journal of Theoretical Biology, 379:79–88, 2015.

[34] H. Ohtsuki, J. Y. Wakano, and Y. Kobayashi. Inclusive fitness analysis of cumulative cultural evolution in an island-structured population. Theoretical Population Biology, 2017.

[35] Y. Kobayashi, J. Y. Wakano, and H. Ohtsuki. Evolution of cumulative culture for niche construction. Journal of Theoretical Biology, 472:67–76, 2019.

[36] C. Mullon, J. Y. Wakano, and H. Ohtsuki. Coevolutionary dynamics of genetic traits and their long-term extended effects under non-random interactions. Journal of Theoretical Biology, 525: 110750, 2021.

